# A gene duplication of a septin provides a developmentally-regulated filament length control mechanism

**DOI:** 10.1101/2021.04.23.441157

**Authors:** Kevin S. Cannon, Jose M. Vargas-Muniz, Neil Billington, Ian Seim, Joanne Ekena, James Sellers, Peter Philippsen, Amy. S. Gladfelter

## Abstract

Septins are a family of conserved filament-forming proteins that function in a variety of processes including cell cycle progression, cell morphogenesis and autophagy. Despite their conservation from yeast to humans, the number of septin genes within an organism varies and higher eukaryotes express many septin isoforms due to alternative splicing. It is unclear how variability in septin complex composition influences the biophysical properties of septin polymers. Here we report that a complex duplication event within the *CDC11* locus in the fungus, *Ashbya gossypii*, gave rise to two similar, but distinct Cdc11 proteins, Cdc11a and Cdc1b. *CDC11b* transcription is developmentally regulated producing different ratios of Cdc11a and b complexes during *Ashbya’s* lifecycle. Moreover, deletion of either *CDC11a* or *CDC11b* results in distinct cell polarity defects. Remarkably, despite substantial identity in amino acid sequence, Cdc11a and Cdc11b complexes have distinct biophysical properties with clear filament length and membrane-binding ability differences. Thus, septin subunit composition has functional consequences for filament properties and such functional plasticity can be exploited for distinct biophysical properties and cell functions.

## Introduction

Septins are a family of filament-forming, GTP-binding proteins that function in many cell processes including cytokinesis (Hartwell *et al*., 1974), cell polarity (Gladfelter *et al*., 2005), and membrane organization (Luedeke *et al*., 2005; Yamada *et al*., 2016). Although septins are highly conserved from yeast through humans, the number of septin genes between organisms can vary greatly, from 1 in *Chlamydomonas* to 13 humans (Pan, Malmberg and Momany, 2007). This variable number of septin genes within organisms has been suggested to result from multiple gene duplications. Additionally, alternative splicing in mammalian cells has the potential to produce a wide array of variability in septin gene-products both between different tissues and even within a single cell (Hilary Russell and Hall, 2011; Sellin, Stenmark and Gullberg, 2014). Despite this well appreciated complexity in subunit composition, how the pool of septin proteins available in a cell contributes to septin properties and functions is poorly understood.

Septin proteins self-assemble into hetero-oligomeric complexes that are rod shaped, generally 32 nm long in fungi and are the soluble, minimal subunit of septins (Bertin *et al*., 2008; Bridges *et al*., 2014). These oligomers concentrate on either membrane surfaces or other cytoskeletal networks (in animals) and elongate through annealing such that the terminal subunits of the oligomers interact to create filaments (Field *et al*., 1996; Bertin *et al*., 2010; Bridges *et al*., 2014). These filaments are highly flexible, however their mechanical properties can be modulated through pairing and interacting proteins to form a variety of higher-order structures such as lattices, bundles and rings (Bertin *et al*., 2010; Garcia *et al*., 2011; Ong *et al*., 2014). The process of oligomeric assembly and polymerization has been studied in most detail in the budding yeast *S. cerevisiae*, where 5 mitotic septins are expressed and arranged in the following order: X-Cdc12-Cdc3-Cdc10-Cdc10-Cdc3-Cdc12-X, where the terminal subunit X can be either Cdc11or Shs1 (Garcia *et al*., 2011). Cdc11 and Shs1 impart different properties on complexes and filaments; however, they are only 36 % identical, making it difficult to dissect the molecular basis for the different properties. The interface between terminal subunits is critical to many parameters relevant to septin assembly and could influence properties such as polymerization and fragmentation rates and propensity for bundling and crosslinking.

We set out to address how sequence variations in the terminal subunit can impact septin filament properties by taking advantage of a duplication of the gene encoding *CDC11* in the genome of the filamentous fungus *A. gossypii*. This results in a situation with two versions of Cdc11, namely Cdc11a and Cdc11b, which show 91% sequence identity. Here, we report that Cdc11a and Cdc11b assemble septin structures in *A. gossypii* in a development-specific manner and that deletion of either *CDC11A* or *CDC11B* results in distinct morphological phenotypes. Remarkably this functional specialization arises from small numbers of changes in the primary sequence. The position of Cdc11 at the termini of complexes, which mediate filament elongation, enables these modest residue changes to have substantial impacts on biophysical features of septin filaments. Specifically, different Cdc11 complexes produce septin filaments of distinct lengths, different membrane binding affinities and sizes of higher-order assemblies *in vivo*. Remarkably, a single amino-acid substitution is sufficient to impart the distinct biophysical properties to the the different Cdc11 complexes. This work reveals a how specialization after gene duplication can generate novel biophysical features that can control the size, shape and function of cytoskeletal polymers.

## Results

### Tandem duplication of the *CDC11* locus in the *Ashbya gossypii* lineage

The *Ashbya gossypii* (“*Ashbya*”) genome encodes eight septin genes, all of which have homologs in *S. cerevisiae* (Gattiker *et al*., 2007). Interestingly, in *Ashbya* there are two copies of the *CDC11* gene on two different chromosomes, in contrast the *S. cerevisiae*, which has a single version of Cdc11. This is especially notable because *S. cerevisiae* underwent a whole-genome duplication after divergence from a common ancestor and thus harbors many retained duplicated genes, but not Cdc11 or other septins. *CDC11a* (AER445C) is on chromosome V of *Ashbya* while *CDC11b* (AFR436C) is on chromosome VI. The 411 amino acids of AgCdc11a share 91% identity (95% similarity) to the 408 amino acids of Cdc11b. We predict that these two paralogs originated from an ancestor that had a tandem duplication of the *CDC11* locus (Figure 1A). Based on the current synteny of the loci, after the duplication event in the *Ashbya* ancestor, a DNA double-strand break presumably occurred between the two *CDC11* copies that were repaired through a complex gene fusion event. This duplication was followed by a complex double-strand repair that presumably occurred relatively recently in the *Ashbya gossypii* evolutionary history because the only other sequenced species closely related to *A. gossypii* with a second syntenic copy of *CDC11* is *Ashbya aceri* (Figure 1B). The *A. aceri* Cdc11a protein is 100% identical to Cdc11a from *A. gossypii* Cdc11a while the *A. aceri* Cdc11b shares 95% identity with *A. gossypi*i Cdc11b. *A. aceri* Cdc11b is also more similar to *A. gossypii* Cdc11a (94% identity) than *A. gossypii* Cdc11b (Figure 1C). Thus, a duplication and genome rearrangement has preserved a lineage of closely related *Ashbya* species with an additional Cdc11 protein. We therefore set-out to examine the cellular and biophysical significance of having a second Cdc11 protein of similar but not identical sequence.

**Figure 1.**
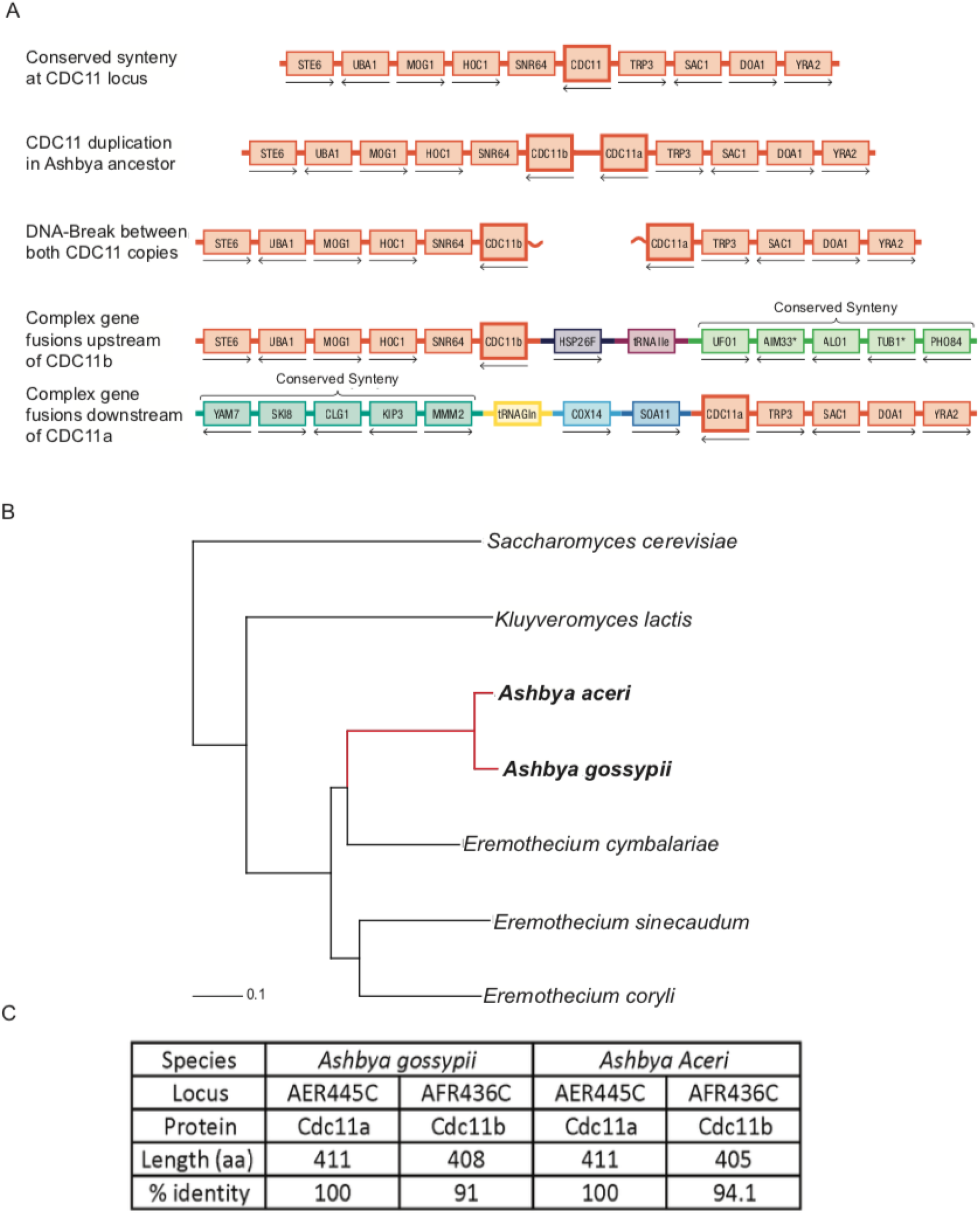
Ashbya genome encodes 2 copies of *CDC11*. (A) Schematics of the molecular origin of *CDC11a* and *CDC11b*. The two copies originated from a tandem duplication followed by a complex double-strand break repair. (B) This duplication occurred only in the *Ashbya* lineage (red line) and both *Ashbya gossypii* and *Ashbya aceri* genomes contain the CDC11b gene. (C) Comparison of the different Cdc11 peptides in *A. gossypii* and *A. aceri*. Percent identity was determined by comparing each peptide to *A. gossypii* Cdc11a.

### Cdc11b is expressed in vivo and colocalizes with Cdc11a

We first examined the expression timing and localization of Cdc11a- and Cdc11b-capped octamers in cells. As the *CDC11b* locus could encode a pseudogene that is not expressed, we examined transcript levels of *CDC11b* over time. We found that both *CDC11a* and *CDC11b* transcripts are regulated throughout the life cycle of *Ashbya*. Specifically, *CDC11b* transcripts are abundant inside spores and following germination, the number of transcripts is reduced until 18 hours, where the *CDC11b* transcript begins to increase in abundance, whereas we see a steady increase in *CDC11a* expression over time through 18 hours (Supplemental Figure S1),

Next, we localized Cdc11 proteins throughout *Ashbya* development. During hyphal growth, septins can assemble into several major structures: 1) Thin filaments: which are dynamic, curving and found all over the cell cortex; 2) Inter-region rings: that consist of bundles of septin filaments aligned parallel to the long axis of the hyphal tube; 3) Branch point structures: where septins localize to sites of micron-scale curvature at the base of emergent hyphae that extend perpendicular to the main hyphal axis; 4) Symmetrical tip-splitting structures: where septins localize to the micron-scale curvature at the bases of bifurcated tips (Helfer and Gladfelter, 2006; DeMay *et al*., 2009; Bridges *et al*., 2016) (Figure 2A). To examine the localization of Cdc11a and Cdc11b-capped octamers at these various structures through time, we generated a strain that co-expressed Cdc11a-mCherry and Cdc11b-eGFP under the control of their native promoters and imaged using confocal microscopy. We observed robust Cdc11a localization at all septin structures from 12 to 24 hours (Figure 2B). The ratio of Cdc11a- to Cdc11b-capped octamer fluorescence intensity at branch point and IR-ring structures shows a consistently higher level of Cdc11a relative to Cdc11b from 12 to 18 hours, which is when all growth is through lateral branches an no tip-splitting has begun (Figure 2C,D). At ∼20 hours, *Ashbya* cease lateral branching and begin to undergo tip-splitting (Knechtle, Dietrich and Philippsen, 2003). Only at 24 hours is Cdc11b localization visible at septin structures and the ratio of Cdc11a- to Cdc11b-capped octamers is reduced (Figure 2C,F,I). However, there is still remnant Cdc11a-capped octamer present at this time at all septin structures, including the tip-split saddle-points, suggesting that these two Cdc11-capped octamers are capable of co-assembling *in vivo*, but the relative abundance of each protein in assemblies depends on the developmental stage of the cells (Figure 2C,F,I).

**Figure 2.**
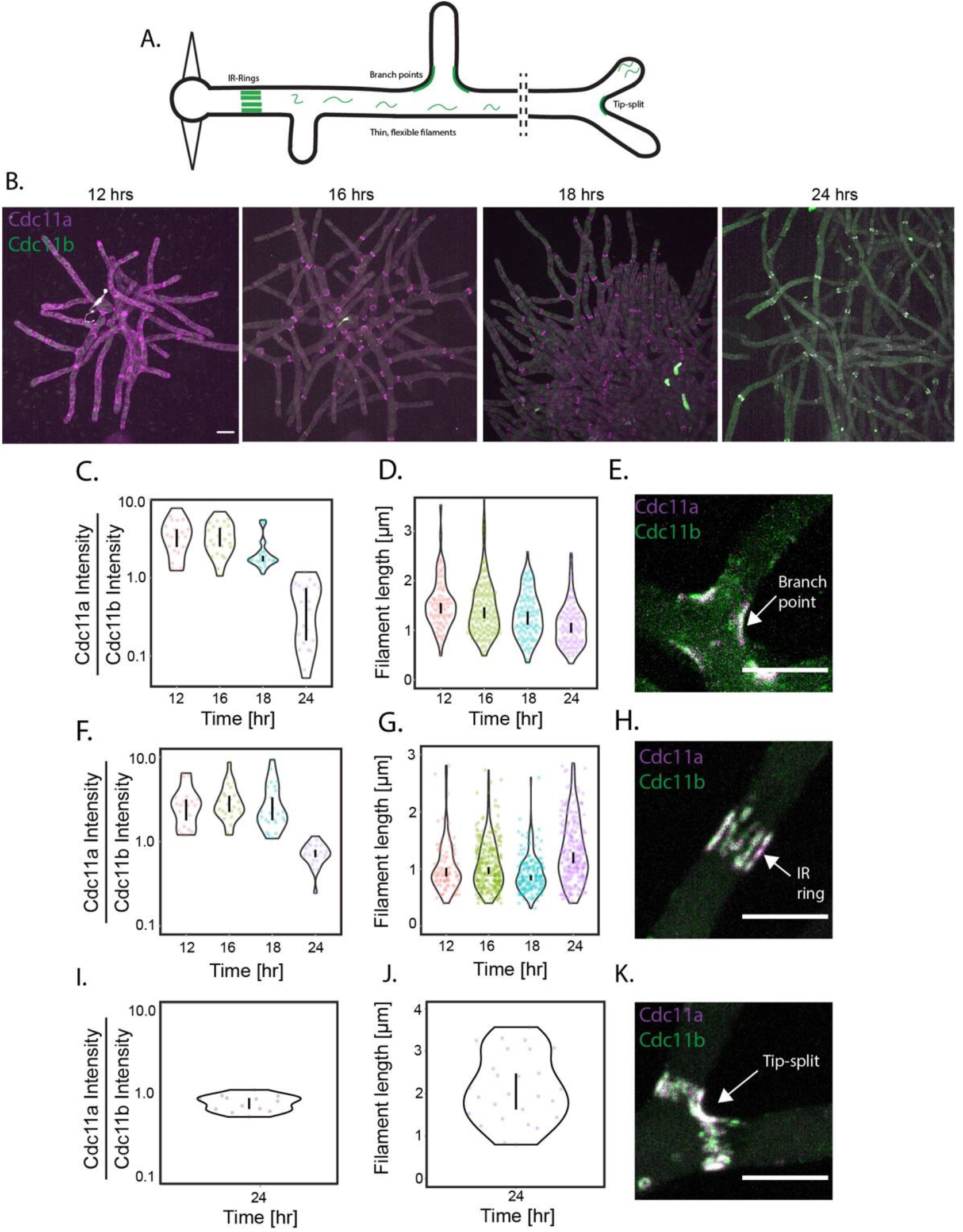
Cdc11a and Cdc11b localization to septin structures is temporally regulated. (A) Cartoon of a simplified *Ashbya* to highlight to variety of septin structure (B) Maximum intensity projections of *Ashbya* cells expressing integrated Cdc11a-mCherry (magenta) and Cdc11b-GFP (green) at different time points. Scale bar 10 µm. (C) Violin plot of the ratio of Cdc11a to Cdc11b sum intensities at branched structures over time. Vertical bars represent the 95% confidence interval. N > 20 septin structures analyzed in > 7 different cells. (D) Violin plot showing measured septin filament lengths at branched structures over time. Vertical bars represent the 95% confidence interval. N > 100 septin filaments analyzed in > 7 cells. (E) Representative maximum intensity projection image of a branch point at 24 hours. Cdc11a (magenta) and Cdc11b (green). Scale bar 20 µm. (F) Violin plot of the ratio of Cdc11a to Cdc11b sum intensities at IR-ring structures over time. Vertical bars represent the 95% confidence interval. N > 20 septin structures analyzed in > 7 cells. (G) Violin plot showing measured septin filament lengths at branched structures over time. Vertical bars represent the 95% confidence interval. N > 100 septin filaments analyzed in > 7 different cells. (H)) Representative maximum intensity projection image of an IR-ring at 24 hours. Cdc11a (magenta) and Cdc11b (green). Scale bar 20 µm. (I) Violin plot of the ratio of Cdc11a to Cdc11b sum intensities at tip-splitting structures over time. N =12 structures cells for N> 10 cells. Vertical bars represent the 95% confidence interval. (J) Violin plot showing measured septin filament lengths at tip-split structures at 24 hours. Vertical bars represent the 95% confidence interval. N > 27 septin filaments analyzed in > 10 cells.

### Cdc11b is associated with changes in higher-order structure size

We next examined if and how the variable stoichiometries of Cdc11a/b influenced the features of higher-order septin assemblies branch points, inter-region rings, and tip-splits. We find that at branch points, the average filament length decreases over time, with the shortest filament occurring at 24 hours, coincident with Cdc11b localization to these structures (Figure 2D,E). Interestingly, in contrast, when we examined septin filament length at IR rings, we found that the average filament length increases over time, with the longest filaments occurring at 24 hours (Figure 2G,H). However, inter-region rings are known to be regulated by the kinases Elm1 and Gin4 (DeMay *et al*., 2009), suggesting that post-translational modification of septins could be another means to regulate filament length in these higher-order structures. Lastly, when we examined septin filament length at tip-splits, we observed that the filaments that follow the curve between the new tips were longer relative to either branch point or IR-ring filaments (Figure 2J,K). Collectively, these data show that Cdc11b expression affects filament length both positively and negatively depending on the type of septin structure.

### Cdc11a and b have distinct functions in cell morphogenesis

Next, we investigated if Cdc11a and/or Cdc11b could functionally compensate for one another. To do this, we generated deletion strains (where either *CDC11A* OR *CDC11B* were deleted from the genome) and examined *A. gossypii* cell morphologies using DIC microscopy. We find that cells lacking *CDC11A* exhibit multiple morphology defects at both 18 and 24 hours, including hyper-branched hyphae (Distance between branches = 21.73 ± 15.5 µm in wild type cells, 13.03 ± 7.73 µm in *cdc11aΔ* cells) and long, persistent hyphae where no lateral branches emerge, as well as aberrant, asymmetric tip-splitting events (Figure 8A,B). Cells lacking *CDC11B* exhibit no observable morphological defects at 18 hours, however at 24 hours we observe aberrant tip-splitting events (Figure 3C) and larger hyphal diameters relative to wild-type (Diameter = 4.45 ± 0.70 µm in wild type cells, 5.75 ± 0.92 µm in *cdc11bΔ* cells). Interestingly, the tip-splitting defects observed in cells lacking *CDC11A* are different than those in cells lacking *CDC11B*. Tip-splitting events hyphae in *cdc11aΔ* are asymmetric and appear to undulate in a wave-like pattern (Figure 8B), whereas *cdc11bΔ* tip-splitting hyphae are either asymmetric or symmetric, and were observed to undergo tri- and quadfurcations (Figure 3C). This is in contrast to wild-type cells where tip-splitting events are always bifurcation events (Figure 3A). This data suggests that Cdc11a and Cdc11b have distinct, developmentally-regulated functions despite their high degree of sequence identity.

**Figure 3.**
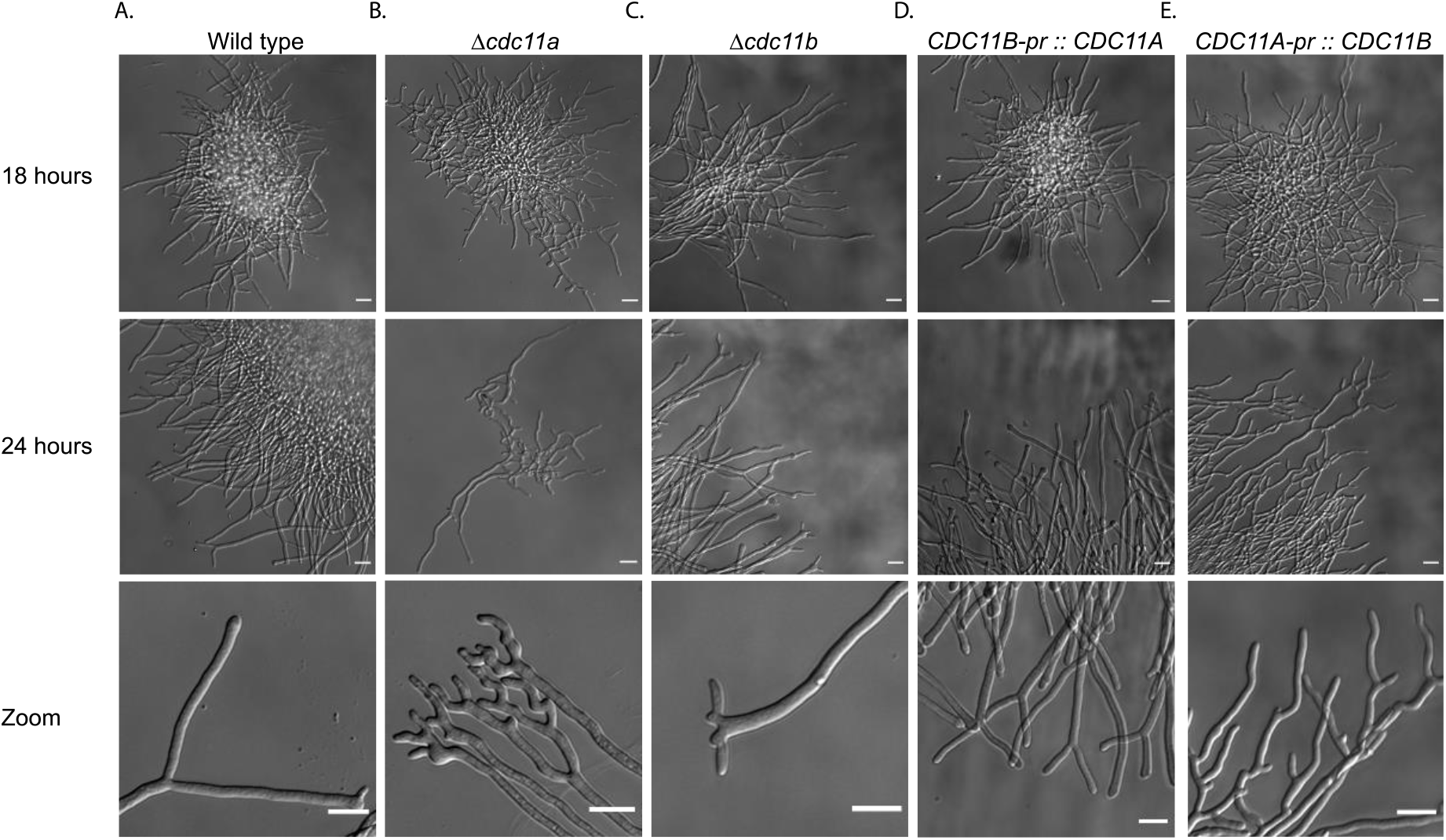
Cdc11a and Cdc11b have distinct morphology depects and cannot fully complement each other. Representative DIC images of *A. gossypii* cells imaged on agar pads after either 18 or 24 hours in liquid culture. Zoomed in image highlights tip-splitting events. All scale bars 20 µm.

To disentangle the degree to which specific functions were due to differences in expression timing or differences in the sequence of the two Cdc11 proteins, we generated strains that would either express two copies of *CDC11A* or *CDC11B*. This also ensures that the dose of septin protein is comparable to wild-type cells throughout development unlike the null mutants. For these experiments, Cdc11a is expressed from both the *CDC11a* and *CDC11b* promoter (*CDC11B*-pr: *CDC11A)* and similarly for Cdc11b (*CDC11A-pr: CDC11B)*. In cells with two copies of *CDC11A*, we observe normal growth relative to that of wild-type at 18 hours, however, at 24 hours we observe a high frequency of tip-splitting events, more so relative to wild-type cells (Figure 3D). Interestingly, these tip-splits appear to be morphologically indistinguishable from that of wild-type cells, suggesting that *CDC11A* can functionally compensate for *CDC11B* with respect to hyphal morphology, however it cannot properly compensate in regulating the number of tip-splitting events. Cells with two copies of *CDC11B* are still hyperbranched and retain aberrant tip-splitting phenotypes reminiscent of *cdc11aΔ* cells (Figure 3E). However, the tip-splitting phenotype is somewhat less severe than *cdc11aΔ* cells, suggesting that *CDC11B* can partially functionally compensate for *CDC11A* in tip splitting. Collectively, these data suggest that despite their high sequence similarity, Cdc11a and Cdc11b proteins have discrete functions in septin higher-order assembly and morphogenesis throughout the growth cycle. We next look to understand the biophysical basis for these distinct cell functions.

### Identity of terminal subunit changes the biophysical properties of septin filaments

We set out to determine if Cdc11a or Cdc11b-capped octamers had distinctive biochemical properties such as length, membrane adsorption rate, and rigidity. To do this, we purified recombinant *Ashbya* (from *A. gossypii)* septins expressed in *E. coli*. Septin complexes capped with either AgCdc11a or AgCdc11b form hetero-octamers and are capable of filament formation in solution (Figure 4A,B). Although septins can polymerize in solution *in vitro*, in fungal cells septin filament formation is dependent on septin-membrane interactions (Bridges *et al*., 2014, 2016). We therefore reconstituted septin assembly with recombinant proteins and planar supported lipid bilayers (SLBs) (Figure 4C). The distribution of filament lengths for both Cdc11a and Cdc11b-capped octamers at steady state could be approximated by a single exponential function, consistent with isodesmic polymer growth models (Figure 4D) (Romberg, Simon and Erickson, 2001; Skillman *et al*., 2013; Woods *et al*., 2021). Fitting to a single exponential function also allowed us to calculate an average filament length despite the diffraction limit of the light microscope and for Cdc11a octamers was 0.72 µm (∼23 octamers) and for Cdc11b octamers 0.52 µm (∼16 octamers) (Figure 4D). This suggests that the affinity for “end-on” Cdc11a-Cdc11a interactions is stronger than Cdc11b-Cdc11b interactions and is consistent with the shorter assemblies seen in a subset of Cdc11b-containing higher-order assemblies in cells.

**Figure 4.**
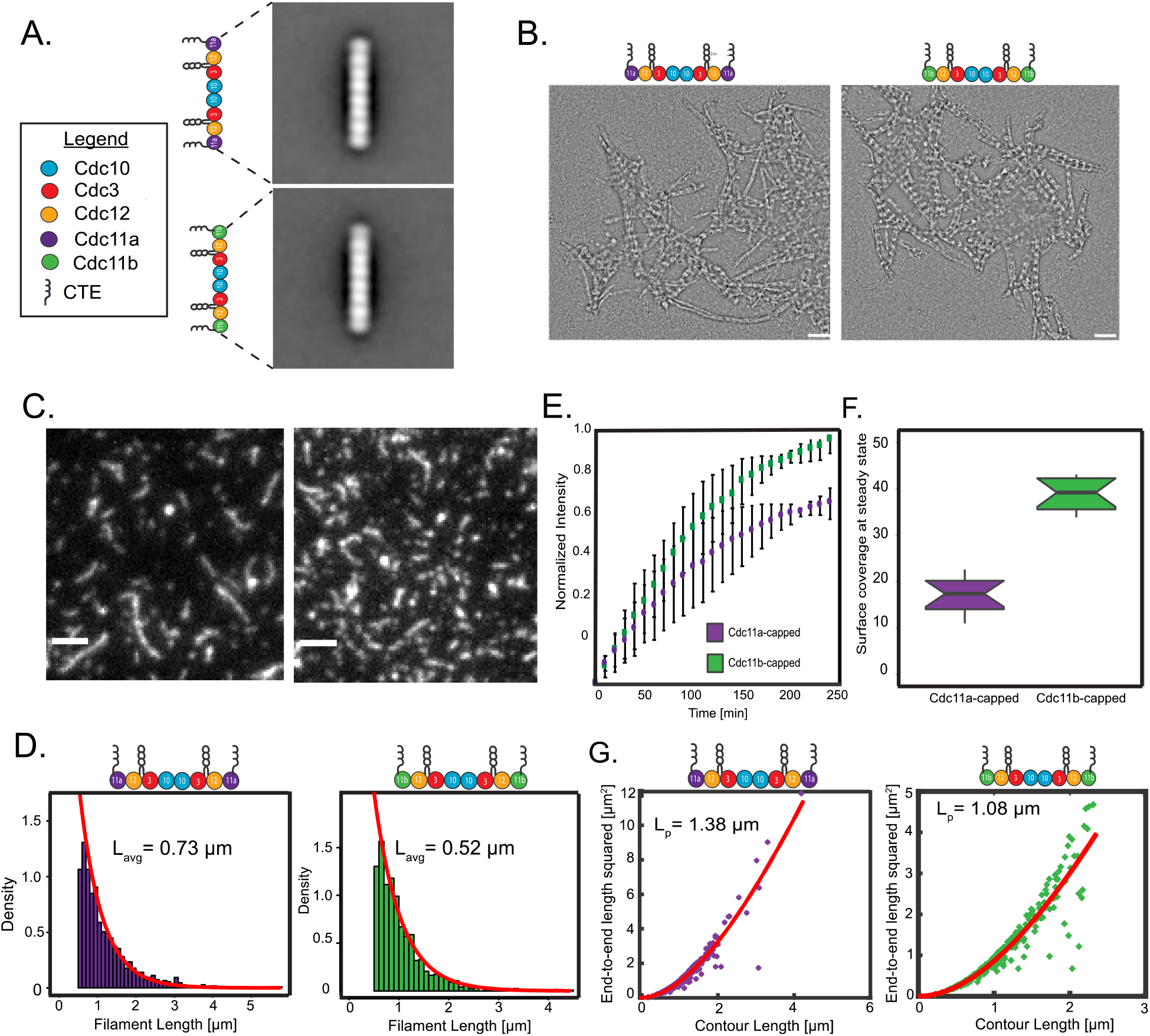
Cdc11a- and Cdc11b-capped octamers have distinct biochemical properties. (A) Global averages and montages of class averages showing the predominant structure of Cdc11a- and Cdc11b-capped octamers in high salt (300 mM KCl). Scale bars, 10 nm Cartoon of septin octamers highlighting individual polypeptides. (B) Transmission electron micrographs of Cdc11a- and Cdc11b-capped octamers forming filaments in solution at low salt (50 mM KCl), respectively. Scale bars, 100nm. (C)Representative TIRF image of 0.5nM Cdc11a- (left panel) or Cdc11b (right panel) -capped filaments formed on a planar supported lipid bilayer (SLBs) (75% DOPC and 25% PI) at steady state. Scale bar 2 µm. (D) Steady-state filament length distributions for Cdc11a- (left panel) and Cdc11b (right panel)- capped octamers. Distributions were fit to a single exponential (red) from which the average filament length was determined. (E) 1nM of either Cdc11a- or Cdc11b-capped octamers were seeded onto planar SLBS (75% DOPC and 25% PI) and imaged by TIRF microscopy through steady state. Normalized fluorescence intensity is plotted as a function of time relative to Cdc11b-capped octamer intensity. (F) Violin plot showing the percent surface coverage for Cdc11a- and Cdc11b-capped filaments at steady state on planar SLBs. Horizontal black bars represent the mean, vertical black bars represent the deviation from the mean. N=10 different locations on the bilayer over three bilayers. (G) End-to end distance analysis was used to calculate the persistence length of Cdc11a-(left panel) and Cdc11b- (right panel) capped filaments. Filled black dots represent individual filaments. Redlines represent the fit. N > 80 filaments for each sample.

We hypothesized that differences in affinity between neighboring Cdc11-capped complexes could influence the rate at which septins assemble onto the membrane, potentially by impacting the off-rate since filament length controls dissociation. To test this hypothesis, we adsorbed septin containing either Cdc11a- or Cdc11b-capped septin octamers onto planar supported lipid bilayers and monitored change in fluorescence intensity over time to measure rates of assembly. Interestingly, we observed similar increase in fluorescence intensity through time for both Cdc11a- and Cdc11b-capped octamers, suggesting septin assembly rates on the membrane are not dependent on the Cdc11 polypeptide that occupies the terminal position within the hetero-octamer (Figure 4E). Surprisingly, however, when we examined the surface coverage for both types of filaments at steady-state, we found that Cdc11b-capped filaments occupy approximately 2-fold greater space on the membrane (Figure 4F). We speculate that the observed higher surface coverage by Cdc11b-capped filaments could result from a shorter average filament length, as shorter filaments might be able to more efficiently pack together on the membrane than longer filaments. Additionally, having a higher density of bound septins with shorter filaments would increase the number of available binding sites (free filament ends) for which septin octamers coming from the bulk could interact, thereby potentially increasing amount of septin bound to the membrane.

The mechanical properties of cytoskeletal filaments are intimately tied to their functions. As Cdc11a or Cdc11b are the terminal subunits for filament formation, they are uniquely poised to influence the intrinsic flexibility of a septin filament. Therefore, next we characterized the persistence length (Graham *et al*., 2014), the average length over which filaments stay straight, for septin filaments composed of either Cdc11a- and Cdc11b-capped octamers. Interestingly, the persistence length of both classes of filaments is similar (1.38 and 1.13 µm, Cdc11a and Cdc11b complexes, respectively) (Figure 4G), suggesting that despite apparently different affinities, that Cdc11a and Cdc11b generate filaments with similar flexibilities. When combined these data show that the identity of septin protein in the terminal position of the oligomer can produce septins with different average lengths and packing densities on membranes, indicating that relatively small changes in the protein sequence can produce substantially different polymer behaviors.

### Cdc11a and Cdc11b septin complexes display the same curvature preference

Septins are the only identified sensor of positive micron-scale membrane curvature in eukaryotes. Previous work in our lab has utilized an in vitro reconstitution system to measure septin adsorption using positively curved SLBs, fluorescently-tagged, recombinant protein, and quantitative fluorescence microscopy (Bridges *et al*., 2016). Recombinant septins from budding yeast were shown to preferentially assemble onto spherical SLBs with a curvature, κ= 2 µm^-1^ (corresponding to a diameter of 1 µm) when mixed with SLBs of various curvatures. We used this spherical SLB system to examine the curvature-sensing ability of *Ashbya* septins containing either Cdc11a or Cdc11b-capped septin octamers. When we measured adsorption for both octamer types at steady state, we found that both Cdc11a and Cdc11b have preference for beads where κ=2µm^-1^ (Figure 5A,B), consistent with the curvature preference for budding yeast septins. Interestingly, we noticed that Cdc11b-capped octamers show a higher absorption value, (∼0.8 and ∼1.0 for Cdc11a and Cdc11b, respectively) (Figure 5B), indicating that more of these septins are bound to the bead. This is consistent with our previous data where Cdc11b-capped octamers occupy a higher surface coverage on planar membranes relative to Cdc11a at steady state (Figure 4E). Higher absorption values onto beads (κ =2µm^-1^) at steady state could be explained through at least two hypotheses. The first being that Cdc11b-capped complexes have a stronger affinity (K_d_) and/or B_max_ for curved membranes than Cdc11a-capped octamers. The second possibility is that Cdc11b capped filaments are packed more efficiently than Cdc11a-capped filaments on these membranes. Unfortunately, we cannot easily distinguish between these scenarios because due to the resolution limit on our confocal microscope (as we cannot resolve septin filaments), combined with low protein yield during purification such that we were unable to generate the saturation binding isotherms to calculate the precise K_d_ and maximum binding values.

**Figure 5.**
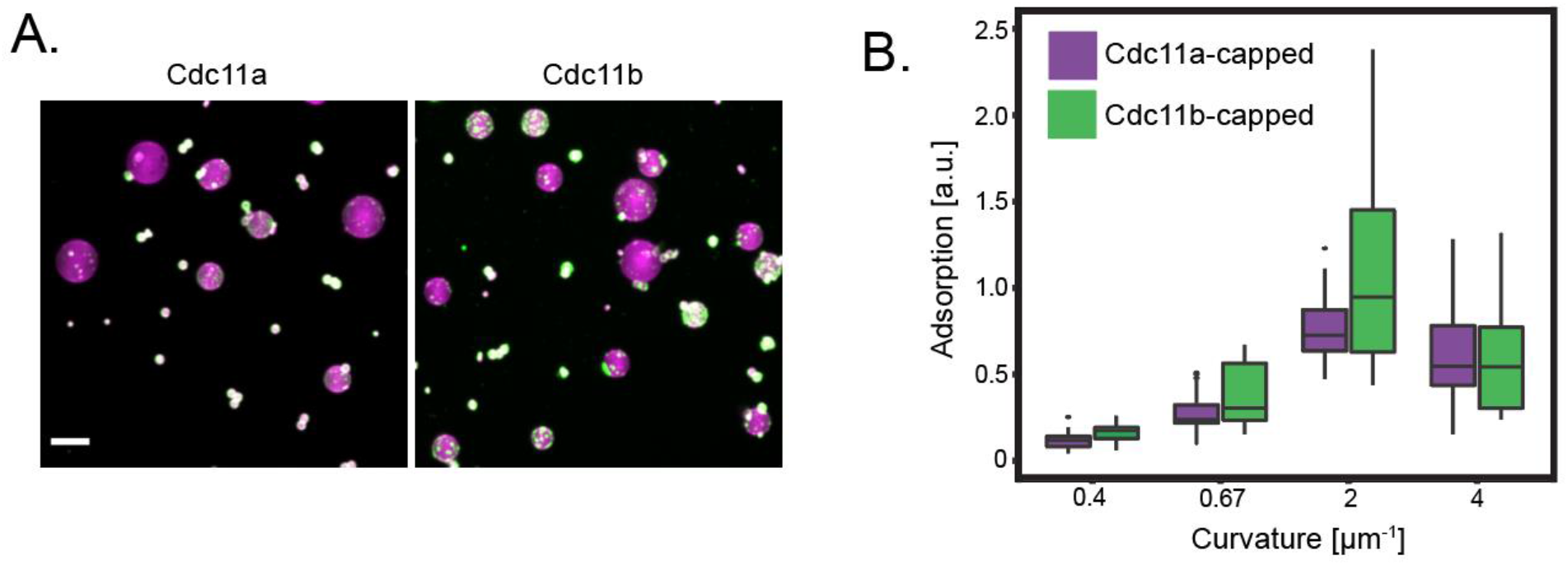
Cdc11a- and Cdc11b-capped filaments display the same curvature preference. (A) SLBS (75% DOPC, 25% PI, 0.05% Rh-PE) were formed onto silica microspheres with different curvatures κ=0.4, 0.67, 2, and 4 µm^-1^). 15 nM of Cdc11a-or Cdc11b-capped septin octamers were added to the microspheres and images were collected at steady state (1hr). Representative images are maximum intensity projections. Magenta (lipid), green (septin). Scale bar 5 µm. (B) Box and whisker plot quantifying septin adsorption onto silica microspheres n>50 beads for each curvature.

### Cdc11a and Cdc11b complexes co-assemble into filaments on planar and curved membranes

As Cdc11a and Cdc11b appear co-assembled in the same structures in cells, we next investigated if recombinant septin complexes containing either Cdc11a or Cdc11b could co-assemble into filaments *in vitro* in the absence of other cellular factors. First, we mixed equal amounts of Cdc11a- and Cdc11b-capped septin complexes onto planar SLBs and examined filaments using two color TIRF microscopy. We observed filaments that contained both Cdc11a and Cdc11b septin complexes (Figure 6A,B), demonstrating that both classes of septin complexes are capable of co-assembling into filaments. However, we did notice that there appeared to be a higher density of Cdc11b-capped complexes on the membrane, despite adding equal concentrations of both septin octamers which was consistent with our previous results (Figure 4E). Interestingly, when we measured the steady-state filament length distribution of co-assembled polymers, we found that the average filament length was shorter than either Cdc11a or Cdc11b filaments alone (Figure 6C, and Figure 4A,B). Moreover, we observed an increase in filament persistence length (from ∼1µm to ∼2µm), suggesting that a “mismatch” interaction between Cdc11a-Cdc11b can influence filament rigidity (Figure 6D). Collectively, these data show that co-assembly of these different septin complexes can lead to polymers of distinct length and flexibility compared to either type of homopolymer.

**Figure 6.**
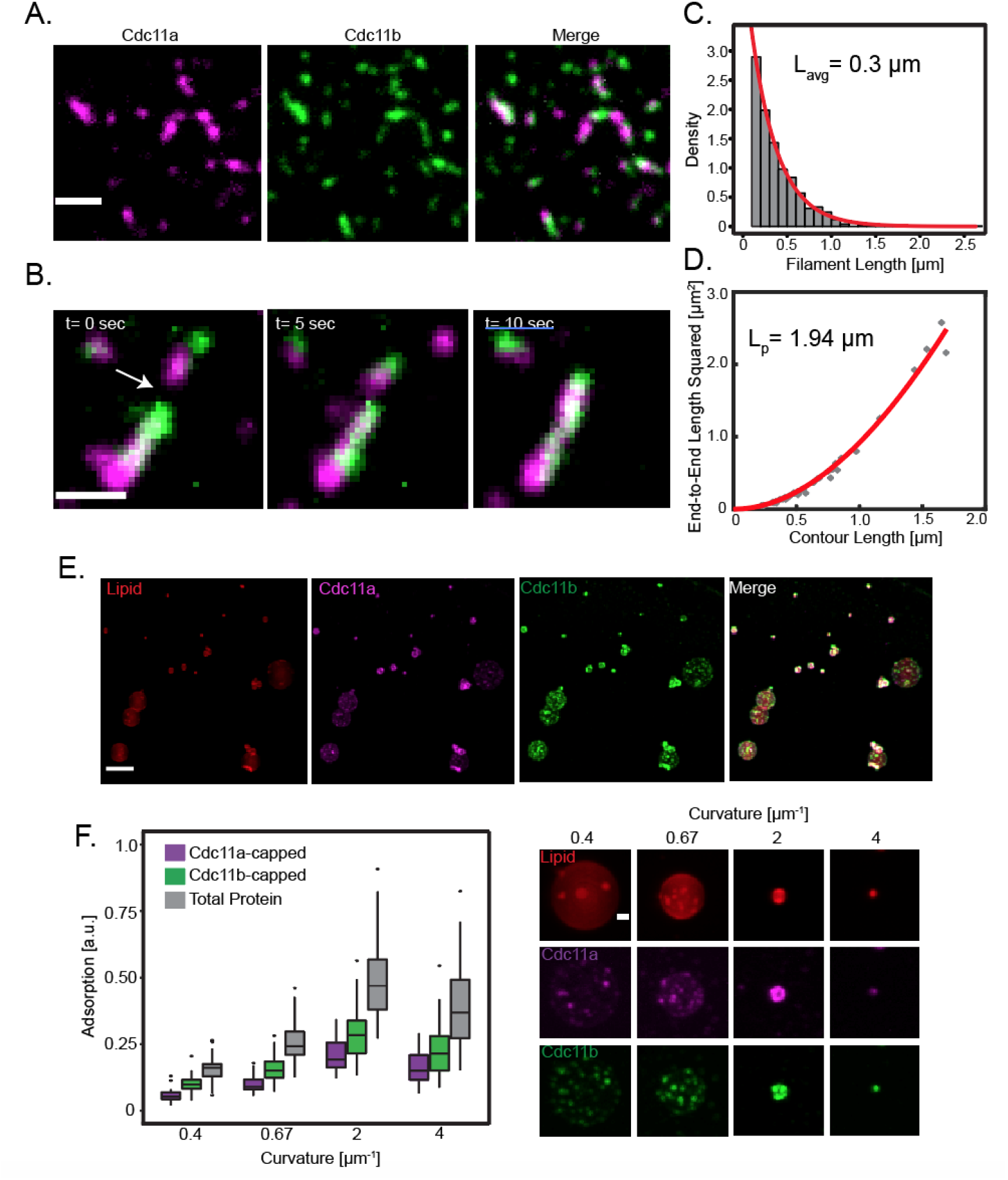
Cdc11a- and Cdc11b-capped octamers co-assemble into filaments on planar and curved membranes. (A) Representative TIRF image of filaments assembled from 0.25 nM of Cdc11a- (magenta) and Cdc11b- (green) capped octamers (total 0.5 nM protein) on planar SLBs at steady state. Scale bar 5 µm. (B) Image montage highlighting an annealing event. Scale bar 1 µm. (C) Steady state filament length distribution. Red line is a single exponential fit from which the average filament length was determined. (D) End-to-end distance analysis was used to calculate the persistence length of co-assembled filaments. Dilled black dots represent individual filaments. Red lines represent the best fit N >60 filaments. (E) SLBS (75% DOPC, 25% PI, 0.05% Rh-PE) were formed onto silica microspheres with different curvatures (κ= 0.4, 0.67, 2, and 4 µm^-1^). 7.5 nM of Cdc11a- and Cdc11b-capped septin octamers (15 nM total protein) were added to the microspheres and images were collected at steady state (1 hour). Representative images are maximum intensity projections. (Red (lipid), magenta (Cdc11a-capped octamers), green (Cdc11b-capped octamers). Scale bar 5 µm. (E) Box and whisker plot quantifying septin adsorption onto silica microspheres. Cdc11a-capped (purple), Cdc11b-capped (green), total protein (grey). Horizontal black bars represent the median and the vertical black bars represent the spread of the data. N >30 beads for each curvature. Next to the plot are representative maximum intensity projections of single beads of various curvature; lipid (red), Cdc11a-capped (magenta), Cdc11b-capped (green). Scale bar 1 µm.

Next, we measured septin adsorption onto SLBs of various curvatures after mixing equal concentrations of Cdc11a and Cdc11b together (Figure 6E,F). Unsurprisingly, we saw a curvature preference for beads where κ = 2µm^-1^ for both Cdc11a- and Cdc11b-capped complexes, consistent with our previous observations (Figure 5A,B). Moreover, Cdc11b complexes showed a higher adsorption than Cdc11a-capped octamers (Figure 6F). Interestingly, we observed less total adsorption of protein onto all tested curvatures when we mixed Cdc11a and Cdc11b-capped octamers (Figure 6F) than we saw with either type of octamer alone (Figure 5B).

### C-terminal extension chimeras do not phenocopy wild-type Cdc11a or Cdc11b filament or curvature sensing properties

Much of the sequence variation between Cdc11a and Cdc11b occurs within the C-terminal extension (CTE) region of the protein (Supplemental Figure S1). Therefore, we generated Cdc11 chimeras through swapping the CTE’s of Cdc11a and Cdc11b to test if the CTE sequence was sufficient to impart the properties particular to either Cdc11a or Cdc11b. Using planar SLBs, we found that Cdc11a-Cdc11bCTE and Cdc11b-Cdc11aCTE-capped octamers had an average steady state filament length of 0.94 µm (∼29 octamers) and 0.55 µm (∼17 octamers), respectively (Figure 7B,D). Interestingly, Cdc11a-Cdc11bCTE-capped octamers are longer than Cdc11a-capped octamers, whereas there is no difference in average filament length between Cdc11b-Cdc11aCTE- and Cdc11b-capped septin octamers. Most surprisingly, we observed equal surface coverage for both types of polymers, despite their unequal average filament lengths (Figure 7C). Additionally, we observed that swapping the CTE’s does not alter filament rigidity, as we calculated similar persistence length values for these complexes (Figure 7E).

**Figure 7.**
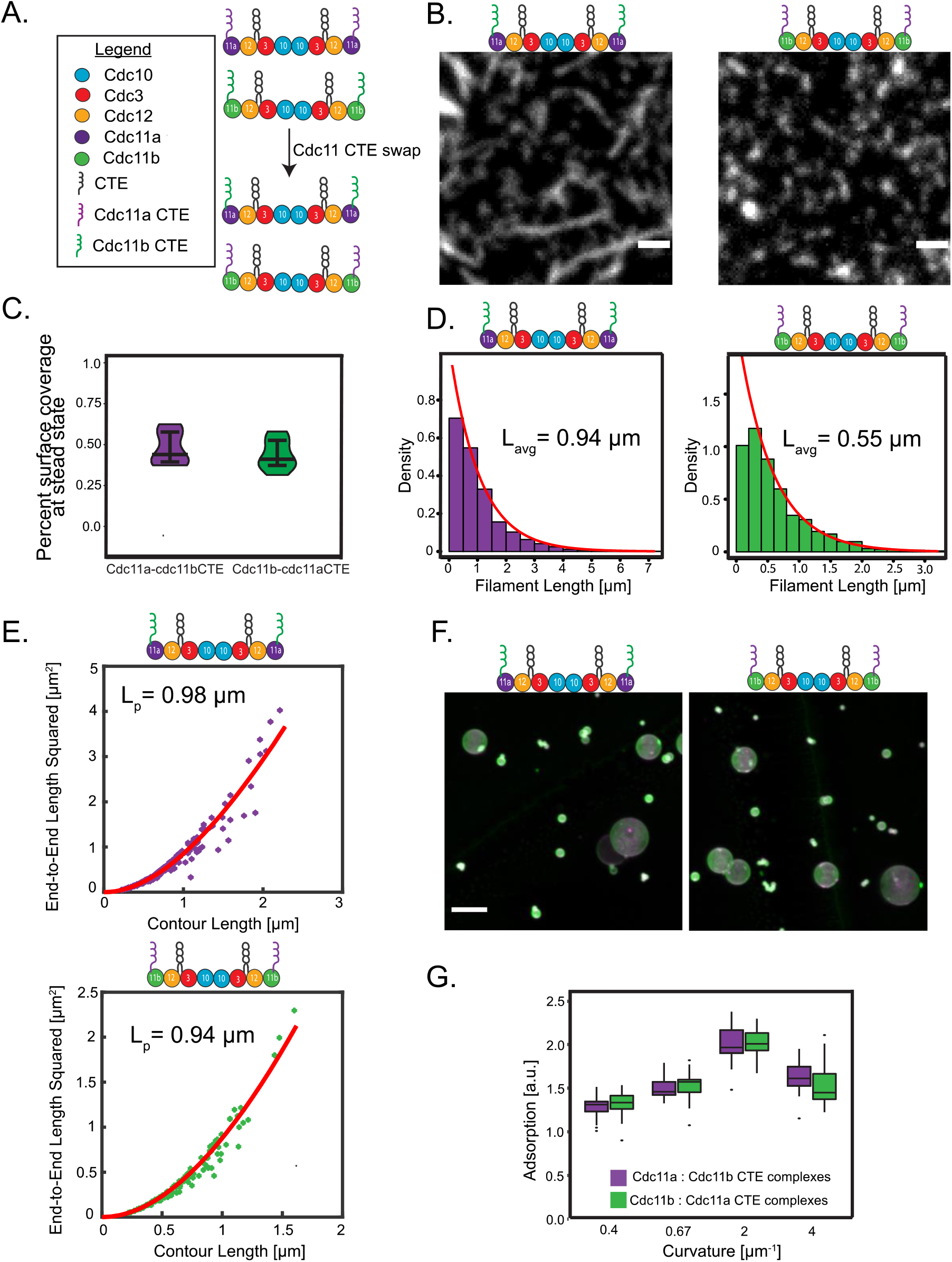
C-terminal extension chimeras do not phenocopy wild-type Cdc11a- or Cdc11b-capped octamer filament or curvature sensing properties. (A) Catoon of septin octamers highlighting CTE swaps. (B) Representative TIRF images of 0.5 nM Cdc11a-Cdc11bCTE (left panel) and Cdc11b-Cdc11aCTE (right panel) capped filaments formed on an SLB (75% DOPC and 25% PI) at steady state. Scale bar 1 µm. (C) Violin plot showing the percent surface coverage for Cdc11a-Cdc11bCTE and Cdc11b-Cdc11aCTE0-capped filaments at steady-state on planar SLBs. Horizontal black bars represent the mean, vertical black bars represent the deviation from the mean. N= five different locations on the bilayer. (D) Steady-state filament length distributions for Cdc11a-Cdc11bCTE (left panel) and Cdc11b-Cdc11aCTE (right panel) capped octamers. Distributions were fit to a single exponential (red) from which the average filament length was determined. (E) End-to-end distance analysis was used to calculate the persistence length of Cdc11a-Cdc11bCTE- (top panel) and Cdc11b-Cdc11aCTE- (bottom panel) capped filaments. N>90 filaments for each sample. (F) SLBs (75% DOPC, 25% PI, 0.05% Rh-PE) were formed on silica microspheres with different curvatures (κ= 0.4, 0.67, 2, and 4 µm^-1^). 15 nM of Cdc11a-Cdc11bCTE- and Cdc11b-Cdc11aCTE-capped octamers were added to the microspheres and images were collected at steady-state (1 hr). Representative imges are maximum intensity projections. Magenta (lipid), green (septin). Scale bar 5 µm. (G) Box and whisker plot quanitfying septin adsorption onto silica microspheres. Horizontal black bars represent the median. Vertical black bars represent the spread of the data. N> 30 beads for each curvature.

Next, we examined if swapping the CTE’s would result in any differences in curvature sensitivity. We found that both chimeras still showed a curvature preference for 1 µm beads (Figure 7F,G). However, we did notice an increase in total adsorption onto the beads for both chimeras relative to their wild-type Cdc11a and Cdc11b-capped octamers (Figure 7F,G). Collectively, these data show that despite the CTE sequences harboring most of the sequence variation between the two Cdc11 paralogs, they are insufficient to recapitulate the biochemical/biophysical features of each Cdc11 polypeptide.

### A single point mutation can tune the filament length distribution of Cdc11a-capped complexes

Given that the CTEs were not sufficient to interconvert the function of the different Cdc11 proteins, we examined the predicted structure of the globular portion of the protein to search for any differences between these two proteins. We threaded both Cdc11a and Cdc11b through Phyre 2 (L. A. Kelley *et al*., 2015) and overlaid the predicted structures (Figure 8A). Despite most of the globular core being highly conserved among both gene products, we found a single sequence variation located just before a prominent difference in predicted structure. Specifically, Cdc11a contains a threonine at position 62, whereas Cdc11b contains an alanine. Interestingly, the Cdc11a sequence flanking this substitution is predicted to be more helical, whereas the structure within this region of the Cdc11b polypeptide is predicted to be disordered. With these potential structural differences in mind, we purified a Cdc11aT62A-capped complexes and examined the steady-state filament length distribution on planar SLBs. Remarkably, we observed that the average filament length of these complexes was dramatically reduced when compared to wild-type Cdc11a-capped octamers, and much closer to the average filament length of wild-type Cdc11b-capped octamers (Figure 8B,C). Moreover, the surface coverage of Cdc11aT62A-capped octamers was similar to Cdc11b-capped octamers (Figure 8D). Interestingly, this portion of the polypeptide is not located at the surface on the canonical polymerization interface (N-C interface). However, as Cdc11 polypeptides are mirror images of one another, we speculate that when neighboring Cdc11-capped octamers oligomerize, this region of the protein could participate in interactions that drive polymerization. Alternatively, by altering this residue, we’ve induced a conformational change at the polymerization interface. In sum, these data show that even a single amino acid difference can alter the degree to which septins form filaments, which is a fundamental feature of the septin cytoskeleton.

**Figure 8.**
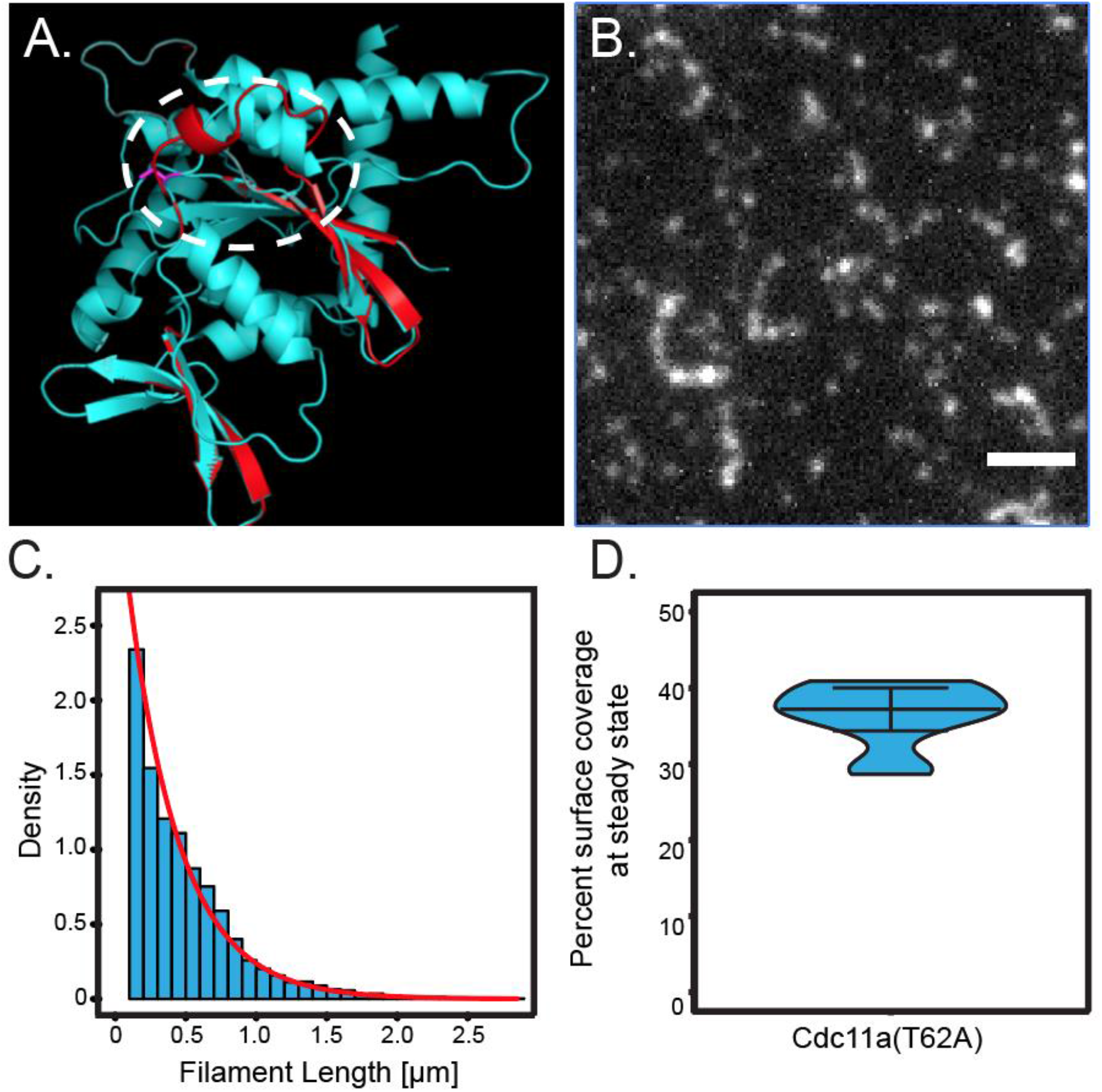
A single point mutation within a novel polymerization interface can tune the filament length distribution of Cdc11a-capped complexes. (A) Cdc11a(cyan) and Cdc11b (red) primary sequences were threaded through the structural prediction algorithm Phyre and overlayed using Pymol. The site of structural divergence is circled and is preceded by a single amino acid difference (magenta). (B) Representative TIRF image of 0.5 nM Cdc11aT62A-capped octamers forming filaments on a planar supported lipid bilayer (75% DOPC and 25% PI) at steady-state. Scale bar 2 µm. (C) Steady-state filament length distriubtuions for Cdc11aT62A-capped octamers. Distributions were fit to a single exponential (red) from which the average filament length was determined. (D) Violin plot showing the percent surface coverage for Cdc11aT62A-capped octamers at steady-state on planr SLBs. Horizontal black bars represent the mean, vertical black bars represent the deviation fro the mean. N= 10 different locations on a bilayer for 3 different bilayers.

## Discussion

In this study we investigated the effects of having two similar, but distinct terminal subunits (Cdc11a and Cdc11b) on septin filament properties using in vitro reconstitution and in vivo imaging. Our data demonstrates that small changes in individual septin subunits can lead to distinct roles in morphogenesis and higher-order assembly sizes in cells and alter septin filament lengths, which in turn can influence the kinetics of septin assembly onto membranes. Collectively, our data highlight how cells might utilize different pools of available septin polypeptides to alter their biochemical/biophysical properties of the septin cytoskeleton to best suit downstream functions.

All known fungi with budding yeast-like genomes, even those which underwent a genome duplication event carry a single copy for each septin. Interestingly, *Ashbya* species are the exception by carrying two copies of *CDC11* (Gattiker *et al*., 2007). What then makes *Ashbya* unique? *Ashbya* has evolved a lifestyle of a filamentous fungus with continuously growing and branching hyphae. The closely related *E. cymbalariae, E. sinecaudum* and *E. coryli* carry one *CDC11* gene, yet also form hyphae. However, none of these three species exhibit two branching patterns, namely lateral branching (low speed growth, 0.2 µm/min for emergent germling hyphae) and symmetrical tip-splitting events (faster growth, 3.5 µm/min within 24 hours) (Köhli *et al*., 2008). It is important to note that lateral branches are the predominant mode of growth early in *Ashbya* development (Knechtle, Dietrich and Philippsen, 2003), whereas tip-splitting events begin later stages of growth (around ∼20 hours) (Ayad-Durieux *et al*., 2000). The genome rearrangements leading to two *CDC11* copies did not involve upstream sequences of *CDC11A*, however, the promotor region of *CDC11B* underwent a rearrangement potentially affecting its regulation. Indeed we see that *CDC11A* is expressed at all developmental growth stages, whereas the expression of *CDC11B* is restricted to the fast tip-splitting phase. This is consistent with our deletion and replacement data that suggests Cdc11a is important in lateral branching and tip-splitting. In contrast, Cdc11b plays a minor role in lateral branching, and is more important later in development for tip-splitting events. Interestingly, cells with two copies of *CDC11B* still show asymmetric tip-splitting events, suggesting that *CDC11A* and *CDC11B* have distinct functions at these sites.

How do septins regulate tip-splitting? This is a fascinating morphological transition that is in major contrast to budding yeast where there is strict singularity in polarization sites, with competition leading to a single, winning site (Howell *et al*., 2009). We speculate that regulating septin filament length/number of septins at tip-splits must be precisely regulated to control the localization and restrict the diffusion of polarity proteins including Cdc42, Bni1, and Spa2 to discrete sites to promote symmetry breaking (Seiler and Plamann, 2003; Schmitz *et al*., 2006; J. B. Kelley *et al*., 2015). Additionally, the morphological defects observed during tip-splitting in either deletion strain could be attributed to the role of septins in cell wall deposition (DeMarini *et al*., 1997) and lipid metabolism (Mela and Momany, 2021) through recruitment of chitin synthases or by the septins themselves providing a mechanical force to counterbalance turgor pressure and cortical tension (Gilden and Krummel, 2010).

Our *in vitro* data show that when Cdc11a- and Cdc11b-capped complexes were seeded onto planar SLBs, Cdc11a-capped complexes formed filaments ∼2x longer than Cdc11b-capped complexes. These data suggest that the affinity between Cdc11a-Cdc11a subunits is stronger than Cdc11b-Cdc11b subunits. However, we were surprised when we saw that Cdc11b-capped complexes showed a higher density of membrane-bound septin than Cdc11a-capped complexes. Our previous data showing that single septin octamers are unable to stay bound (high off rate) to the membrane suggesting that septins must form filaments (to effectively lower the off rate) to remain associated with the membrane (Bridges *et al*., 2016; Cannon *et al*., 2019). Thus, we thought that septin complexes that could form longer filaments would result in a higher density of septins on the membrane. In contrast, we see that Cdc11b-capped complexes, despite forming shorter filaments, show a higher degree of membrane binding than Cdc11a-capped complexes. We speculate that the interaction strength of Cdc11b-Cdc11b interactions is strong enough to form filaments that are capable of staying bound to the membrane (an effective off rate that is low). Furthermore, because the affinity of Cdc11b-Cdc11b interactions is weaker than Cdc11a-Cdc11a interactions, this could allow a higher number of shorter, but stably bound filaments to form on the membrane, effectively increasing the number of binding sites (free filament ends) for which single octamers, either coming from the bulk solution, or diffusing on the membrane could interact. This, in turn, would increase the density of septins on the membrane (Figure 9). Thus, the shorter filaments formed by Cdc11b have essentially a comparable and very low off rate as the filaments formed from Cdc11a, but by being shorter, there are more binding sites at the ends to enhance recruitment of new subunits, yielding a higher density of septins bound to the membrane. However, this might not be due to filament length alone as Cdc11a-Cdc11bCTE-capped complexes show similar membrane binding density despite having different filament lengths. Interestingly, CTE regions of septins has been shown to influence septin filament pairing (Bertin *et al*., 2010). We speculate that these difference is surface coverage may also be due to filament pairing, however we were unable to detect any differences in fluorescence intensity or filament width by light microscopy methods (data not shown). Future work will have to use higher resolution methods to test this hypothesis and pairing may be also relevant in cells.

**Figure 9.**
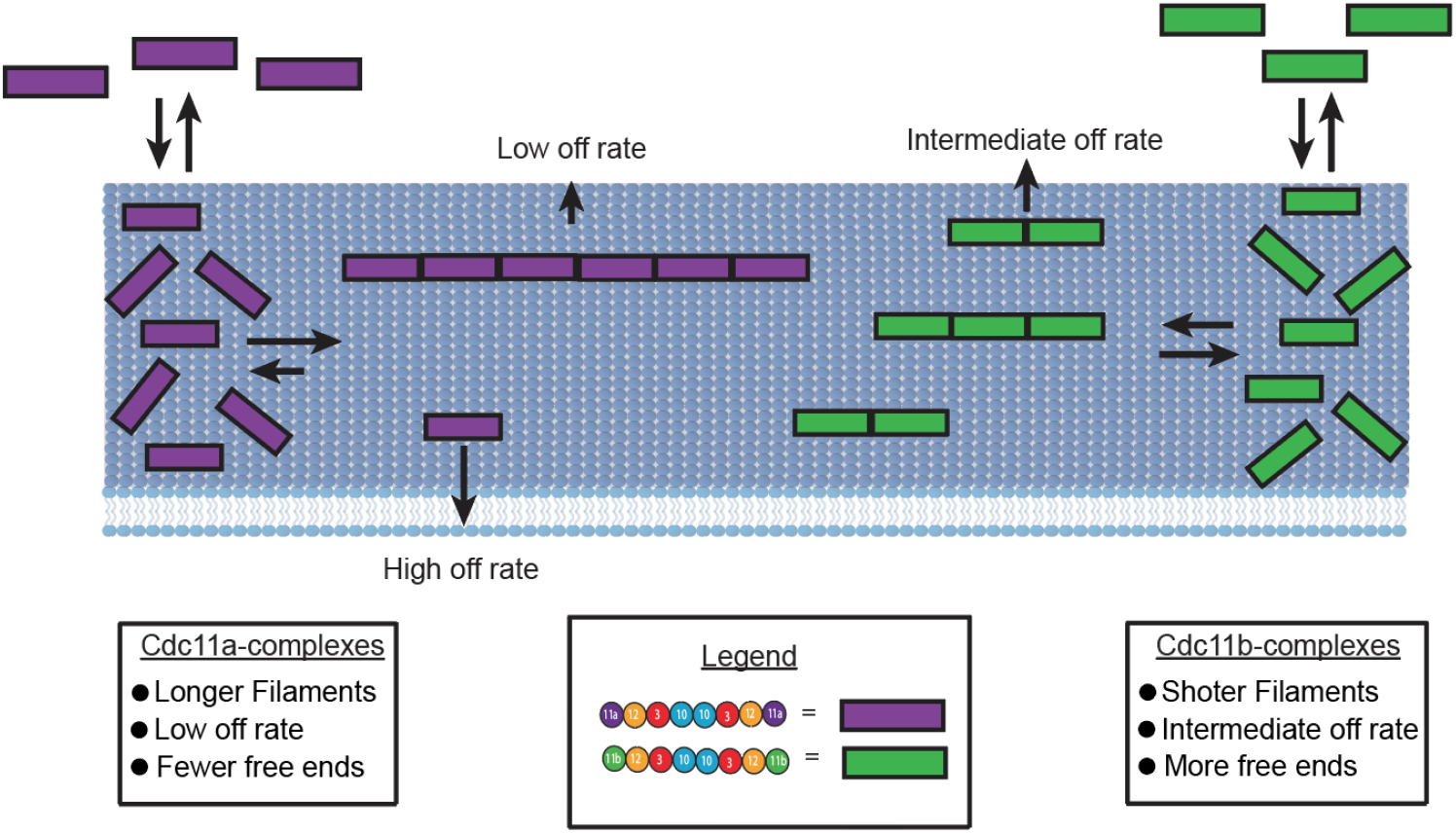
Kinetic argument for how short filaments promote higher membrane adsorption by septins. Single septin octamers from the bulk initially bind to the membrane with the same on and off rates. Through diffusion, septin octamers will collide to form fialments. The overeall length of the filaments depedents on the interaction strength between Cdc11 interactions, such that Cdc11a-capped octamers form longer filaments than Cdc11b-capped octamers. However, not all septin octamers will get incorporated into filaments before dissociating from the membrane (due to the high off rate). Although Cdc11b-capped filaments are shorter than those with Cdc11a caps, there is a greater number of filaments on the membrane. This in turn provides more binding sites (free filament ends) for nearby octamers on the membrane / in bulk solution to interact leading to an increase in adsorption of Cdc11b-capped octamers.

Interestingly, we found that Cdc11a- and Cdc11b-capped octamers are capable of co-polymerizing into filaments that show different biochemical/biophysical properties than filaments arising from either type of octamer alone. We saw that co-polymerized filaments showed an increase in filament rigidity relative to filaments capped with either complex alone, highlighting the importance of the terminal subunit in controlling filament flexibility. It should be noted, however, that the persistence length values reported here are lower than those previously reported for septins. However, we are confident in our values as we have measured a higher number of filaments than in our previous reports ((Bridges *et al*., 2014; Khan, Newby and Gladfelter, 2018) and used both the end-to-end and cosine correlation analysis (data not shown) methods to calculate persistence length (Graham *et al*., 2014), which yielded similar values.

When Cdc11a- and Cdc11b-capped complexes are mixed at a 1:1 ratio the average filament length at steady-state and total adsorption onto membrane curvature is decreased relative to Cdc11a or Cdc11b-capped complexes. Consistent with this, the average filament length at branch points decreases as Cdc11b expression increases. In contrast, as Cdc11b expression increases, we observe longer filaments at IR-rings. However, IR-rings are known to be regulated by the kinase Elm1 (DeMay *et al*., 2009), suggesting that septin filament length can be further regulated by post-translational modifications. As a major function of septins is to serve as a scaffold, we suspect that by controlling the local concentration of septins (through regulating filament formation) at the membrane, the cell can subsequently tune the concentrations of septin-interacting proteins that promote processes such as polarized growth, cell wall deposition, and mitotic progression. Future work should determine the precise stoichiometry between Cdc11a- and Cdc11b-capped octamers and how these relative levels regulate the total septin concentration on the membrane.

The septin superfamily is thought to have emerged from ancient gene duplications that gave rise to the different classes of septins shared across many eukaryotes (Pan, Malmberg and Momany, 2007; Auxier *et al*., 2019). This recent duplication of a terminal septin subunit reveals how small changes in the primary sequence can lead to substantial biophysical differences for the polymer providing a glimpse at how functional diversification may have occurred in this gene family.

## Materials and Methods

### Plasmid construction

Plasmid AGB141 was generated by PCR amplification of a fragment containing mCherry-NAT using AGO190/AGO299 from AGB048 (pAGT211) and integrating it into AGB125 via yeast homologous recombination. Plasmid AGB1320 was generated by PCR amplification of a fragment containing GFP:GEN using AGO2823/AGO2824 from AGB005 (pG3GFP) and integrating it into AGB124 via yeast homologous recombination. Plasmids AGB1369 (Cdc11A::G418) and AGB1370 (Cdc11B::G418) were both generated via yeast homologous recombination. The G418 resistance cassette was PCR amplified using either AGO3326/AGO3327 (CDC11A) or AGO3324/AGO3325 (CDC11B) from AGB005 (pG3GFP). The resulting fragment was transformed into Saccharomyces along with EcoNI digested AGB125 (for CDC11A) or SnaBI digested AGB124 (for CDC11B). Plasmids were confirmed by digest and the G418 insert and was sequenced using AGO521 and AGO403. Plasmid AGB1374 (cdc11a::CDC11B-GFP-NEO) was generated via Gibson Assembly using NEB HiFi Assembly Kit in stepwise fashion. First a 3’ fragment of CDC11A was PCR amplified using AGO3336/AGO3337 and cloned into BamHI/XbaI digested pUC19. Second, a fragment containing CDC11B-GFP-G418 was PCR amplified from AGB1320 using AGO3338/AGO3339 and cloned into BamHI digested plasmid generated from the first step. Finally, a 5’ fragment of CDC11A was PCR amplified using AGO3340/AGO3341 and cloned into ApaI digested plasmid resulting from the second step. AGB1352 (cdc11b::CDC11A-GFP-G418) was generated by Gibson Assembly. PCR fragments were amplified using AGO3285/3286 (5’ upstream CDC11B), AGO3287/AGO3288 (CDC11A-GFP-G418) and AGO3289/AGO3290 (3’ upstream CDC11B) from AGB124 (for CDC11B) or AGB214 (for CDC11A-GFP-G418). The three fragments and BamHI digested pUC19 were combined into an assembly reaction. Resulting clones were confirmed by digest and the entire insert was sequenced. All plasmids were confirmed by restriction digest and underwent Sanger sequencing to confirm the fidelity of the target gene sequence.

### Ashbya strain construction

*Ashbya gossypii* strains used in this study can be found in Table S6.1. Description of the plasmids and oligos used to generate the different strains can be found in Table S6.2 and Table S6.3, respectively. To create AG230 (CDC11A-mCherry-NAT), a 4949bp DNA fragment was obtained by digesting AGB141 with HindIII/NheI, electroporated into WT Ashbya gossypii and grown on agar media containing 100ug/ml clonNAT (EMD Millipore). Transformants were verified by PCR and spores were isolated to obtain homokaryons. Strains expressing both CDC11A-mCherry-NAT and CDC11B-GFP-NEO were constructed by transforming AG230 with a 7678bp fragment from PciI/NotI/PspXI digested AGB1320 and selecting on media containing 100ug/ml G418 (CalBiochem). Transformants were verified by PCR. Spores were dissected from multiple isolates to generate homokaryotic strains to be used for microscopy. It was noted that not all homokaryotic spores from an isolate showed GFP fluorescence. Deletions of CDC11A and CDC11B were made by transforming WT Ashbya with a 4480bp fragment or a 4000bp fragment from Xba1/AgeI/NcoI digested AGB1369 and AGB1370 respectively, and growing on G418 selection. Transformants were verified by PCR however no homokaryotic spores were able to be recovered. To make a strain in which the CDC11A ORF was swapped for the CDC11B ORF AGB1374 was digested with ApaI/SbfI, transformed by electroporation into WT Ashbya and grown on G418 selection. To make a strain in which the CDC11B ORF was swapped for the CDC11A ORF a PCR fragment was amplified using template AGB1352 and AGO3285/AGO3290, transformed by electroporation into WT Ashbya and grown on G418 selection. Isolates of both CDC11 swapped strains were verified by PCR. Recovery of homokaryotes was not attempted.

### Generation of Phylogenetic Tree

We utilized Treehouse to generate a subtree of the *Ashbya* lineage (Steenwyk and Rokas, 2019). This tree was based on a previously described Saccharomycotina yeast phylogeny (Shen *et al*., 2018). NCBI Blast was used to identify homologs of *CDC11* in the different species.

### Microarray analyses

Transcription data was obtained from the Ashbya Genome Database. In short, highly purified spores from the ΔlΔt laboratory strain were used and all the data was obtained in duplicate. For the time course experiment, spores were inoculated into AFM broth and samples were acquired at each timepoint. The center represents a region enriched for sporulation, while the rim represents a region of high speed hyphal growth via tip-splitting.

### *Generation of* Ashbya gossypii *septin expression plasmids*

Plasmid AGB1281 was obtained by restriction enzyme-dependent cloning. CDC11b was amplified from AGB124 using primers AGO2823 and AGO2824. AGB119 and PCR products were digested with XhoI and NdeI. Then the digested products were ligated and transformed into *E. coli* DH5-*α*. C terminal extension (CTE) swaps were achieved via fusion PCR followed by restriction enzyme-dependent cloning. Cdc11a N terminus was amplified using AGO3069 and AGO3103, Cdc11a CTE was amplified using AGO3070 and 3102, Cdc11b N-terminus was amplified with AGO2825 and AGO3103, and Cdc11b CTC was amplified using AGO2826 and AGO3102. To generate the Cdc11a-Cdc11bCTE construct, we performed Phusion PCR using AGO2825 and AGO3070. For the Cdc11b-Cdc11aCTE we used the AGO3069 and AGO2826 oligos. AGB and fusion PCR products were digested with NdeI and XhoI. Then ligated and transformed into *E. coli* DH5-alpha. Cdc11a T62A was generated using the Q5-site directed mutagenesis kit (NEB).

### Septin complex purification

Bl21 (DE3) E. coli cells were transformed using a duet construct expression system using ampicillin and chloramphenicol selection (Bridges *et al*., 2014). Selected transformants were then grown overnight in 25 mL of luria broth with ampicillin and chloramphenicol selection at 37 degrees Celsius while shaking. 1/60 of LB liquid cultures were added to 1 liter of terrific broth with ampicillin and chloramphenicol selection and were grown to an O.D.600nm between 0.6 and 0.8. Upon reaching the appropriate O.D.600nm, 1 mM of isopropyl-B-D-1-thiogalactopyranoside was added to the cultures to begin induction. To achieve a stoichiometric septin complex of Ashbya septins, we use Cdc12 with primary sequence derived from *S. cerevisiae* while all other subunits were Ashbya-derived sequences. Induced cultures were grown for 18 hours at 18 degrees Celsius while shaking before harvesting by centrifugation at 10,000 RCF for 15 minutes. Pellets were resuspended in lysis buffer (1M KCl, 50 mM hepes pH 7.4, 40 mM imidazole, 1 mM MgCl2, 10% glycerol, 0.1 % Tween-20, 1mg/ml lysozyme, and a 1x protease inhibitor tablet (Roche) at 4 degrees Celsius for 30 mins to generate cell lysates. Cell lysates were then sonicated on ice for 10 seconds every two minutes. Lysates were clarified by spinning at 20,000 RPM for 30 minutes using an SS-34 rotor at 4 degrees Celsius. The supernatant was passed through a 0.44 um filter and then incubated with 2 mL of equilibrated cobalt resin at 4 degrees for 1 hour. The lysate-resin mixture was then added to a gravity flow column. Following the initial flow-through of unbound lysate, the resin was washed 4x (50 mL each wash) using wash buffer (1M KCl, 50 mM hepes pH 7.4, 40 mM imidazole, 5% glycerol). Bound protein was then eluted using elution buffer (500 mM imidazole, 300 mM KCl, 50mM hepes pH 7.4, 5% glycerol) and then dialyzed into septin storage buffer (300mM KCl, 50 mM hepes pH 7.4, 1mM BME, and 5 % glycerol) for 24 hours using two steps. Protein purity was determined via SDS-PAGE and protein concentration was determined using a Bradford assay.

### Electron microscopy and Image processing

Proteins were diluted to 50 nM (in buffer containing 10 mM MOPS, 2 mM MgCl_2_, 0.1 mM EGTA (pH 7.0) with 50 mM or 300 mM NaCl as indicated in the text. 3 hours after dilution, samples were applied to UV treated, carbon coated EM grids and stained using 1% uranyl acetate. Micrographs were recorded on a JEOL 1200EX microscope using and AMT XR-60 CCD camera at a nominal magnification of 40000x. Rod shaped particles were picked manually for each dataset (n=2300 for CDC11a, n=1777 for CDC11b. Reference-free image alignments and classifications were conducted using SPIDER software. Each aligned dataset was classified into 100 classes using K-means classification and 30 example class averages were selected to produce each montages shown in Figure S1. The whole micrographs shown in Figure S1 are bandpass filtered between 2 and 40 pixels for clarity (FIJI FFT bandpass filter).

### Preparation of small unilamellar vesicles for seeding onto supported lipid bilayers

75 mole percent of dioleoylphosphatidylcholine (DOPC) and 25 mole percent phosphatidylinositol (soy) were mixed in chloroform in a glass cuvette. For bead assays, 0.05 mole percent of rhodamine-phosphotitdyl-ethanolamine (Rh-PE) was added to the above mixture. Lipid films were made by evaporating the chloroform using argon gas followed by an overnight incubation under negative vacuum pressure. The following day, the lipid films were hydrated using an aqueous supported lipid bilayer (SLB) buffer (150 mM KCl, 20 mM hepes pH 7.4, and 1 mM MgCl_2_) to a final concentration of 5 mM. Hydrated lipid films were vortexed for 10 seconds and allowed to sit at 37 degrees Celsius for 5 minutes. This process of vortexing and incubation at 37 degrees Celsius was repeated five more times (or until the lipid film was hydrated). The hydrated film was subject to water bath sonication at 2-minute intervals until the opaque solution became transparent (vindication of small unilamellar vesicle formation).

### Preparation of supported lipid bilayers onto planar and curved surfaces

Planar supported lipid bilayers were prepared by plasma (oxygen) treatment of glass coverslips (PE25-JW, Plasma Etch) on high power for 15 minutes. Reaction chambers were prepared onto the plasma treated glass using a previously established method (Bridges and Gladfelter, 2016). 1 mM of small unilamellar vesicles was added to the reaction chamber (using SLB buffer) followed by the addition of 1 mM CaCl_2_, followed by incubation at 37 degrees Celsius for 20 minutes as to promote bilayer formation. After bilayer formation, excess small unilamellar vesicles were washed vigorously with 150 µL of SLB buffer six times. It is essential not to touch the cover glass during this step as bilayer continuity can be affected. Prior to adding septins to the membrane, the bilayers are washed again with 150 µL of reaction buffer to lower the salt as to promote septin binding (50 mM hepes pH 7.4, 0.13 mg/mL bovine serum albumin (BSA), 1 mM *β*-mercaptoethanol). This step is repeated 5 more times. On the last wash, remove 125 µL of the reaction buffer. 25 µL of septins at the desired concentration were added to the bilayer and imaged by total internal reflection fluorescence microscopy using a Nikon TiE TIRF system equipped with a solid-state laser system (15 mW, Nikon LUn-4), a Nikon Ti-82 stage, a 100x Plan Apo 1.49 NA oil lens, and Prime 95B CMOS camera (Photometrics).

Supported lipid bilayers on silica microspheres were generated as previously reported by (Bridges *et al*., 2016). 50 nM of small unilamellar vesicles (75 mole percent DOPC, 25 mole percent Soy PI, 0.05 mole percent Rh-PE) were added to silica microspheres of various membrane curvatures at a total surface area of 440 mM^2^. Note: The total surface area of each bead size is equal. SUVS were incubated with the microspheres for 1 hour at room temperature on a rotator to induce bilayer formation. After bilayer formation, the microspheres were spun down each bead size at the minimal sedimentation velocity for each bead (It is important not to exceed this number as bilayer continuity can be affected). 50 µL of the supernatant was removed and discarded. 200 µL of pre-reaction buffer (33.3 mM KCl and 50 mM hepes pH 7.4) was used to resuspend / wash the beads. The microspheres were spun down again (at the appropriate sedimentation velocity) and 200 µL of supernatant was removed. 200 µL of fresh pre-reaction buffer was added to resuspend / wash the microspheres. This process was repeated three more times. Washed microspheres were then mixed at a 1:1 ratio. 29 µL of this mixture was added to 721 µL of reaction buffer (33.3 mL KCl, 50 mM hepes pH 7.4, 0.1% methyl cellulose, 0.13 mg/ml BSA, 1 mM *β*-mercaptoethanol). 75 µL of this mixture was added to a reaction chamber glued to a poly-ethylene glycol coated coverslip ((Bridges *et al*., 2016; Cannon *et al*., 2019). 25 µL of septins at a desired concentration were added to the reaction chamber and allowed to reach steady state (1 hr) and were imaged using spinning disc confocal microscopy.

### Measuring protein adsorption on lipid bilayers supported on silica microspheres

Images of fluorescent-tagged septins adsorbed onto curved supported bilayers on microspheres were acquired using a spinning disc (Yokogawa W1) confocal microscope (Nikon Ti-82 stage) using a 100x Plan Apo 1.49 NA oil lens and a Prime 95B CMOS camera (Photometrics). Images were analyzed using Imaris 8.1.2 software (Bitplane AG) as previously described (Bridges *et al*., 2016; Cannon *et al*., 2019).

### Analysis of septin surface coverage onto planar supported lipid bilayers

10 locations on a given lipid bilayer were blindly selected to form 10 separate images. Each image was background subtracted. The total number of pixels with septin signal were divided by the total number of pixels in the image to get the fraction of septins bound to the surface. This number was multiplied by 100 to get a percent surface coverage.

### Filament length distribution measurements and exponential fit analysis

Filament lengths were measured by uploading raw images to FIJI and using the Ridge detection (Steger, 1998) plugin.. After image segmentation and processing, septin length distributions are extracted from each field of view. We are unable to resolve the frequencies of the smallest septin filaments due to the diffraction limit of light (∼200nm) and the small size of septin octamers (∼32nm) and are thus left with an incomplete length distribution. Therefore, an arithmetic average of the observed lengths will not be an accurate estimate of the true mean length of the population, and we must use a model to extrapolate to the true length distribution. Observation, physical models of septin polymerization, and robust model fits suggest that this distribution is a left-truncated exponential. Since we have incomplete data, we utilize a convenient property of exponential distributions to obtain model fits and estimate the population mean length.

An exponential length distribution has the following PDF: *f*(*x*)=*λ*exp(−*λx*), where 1/*λ* is the mean length. If we let *X* be an exponentially distributed random variable, i.e. *X* ∼ Exp(*X*), then *X* is *memoryless*, which means that: *P*(*X*>*x* +*a*|*X*>*a*)=*P*(*X*>*x*), for some cutoff value *a*. In practice, this implies that if the true septin length population is exponentially distributed with some mean length 1/*λ*, then our left-truncated data will be described by the same *λ*. To fit, we simply choose a cutoff value *a* below which data is ignored, subtract this value from the observed lengths, and obtain the fit parameter *λ*. To ensure robustness of fits, fit quality for a small range of cutoff values is assessed, and the smallest cutoff value that corresponds to a stable value of the fit parameter, *λ*, is the one used for the final model fit. Cutoff values are close to 200nm. The true length distribution is related to the left-truncated data by the scaling factor, exp (*λa*).

### Persistence length measurements

Persistence length measurements were calculated from raw TIRF microscopy images of septin filaments seeded onto planar supported lipid bilayers using a previously published MATLAB GUI method (Graham *et al*., 2014)

### Ashbya cell growth and imaging

Ashbya spores were inoculated into full medium at 30 °C for either 12, 16, 18,or 24 hours before harvesting mycelial cells. Harvested cells were washed 3x using low fluorescence media. Cells were subsequently mounted onto low fluorescence medium based agar pads (2% agar). Images were acquired using a spinning disc (Yokogawa) confocal microscopy (Nikon Ti-82 stage), a 100x 1.49 NA oil lens, and a 95B Prime sCMOS camera (Photometrics). Ashbya deletion and replacement strain images were acquired using a wide-field microscope equipped for differential interference contrast imaging. (Nikon Ti-82 stage, a 40x air Plan Apo 0.95 NA objective using a 95B Prime sCMOS camera (Photometrics).

### Branch point, inter-region ring filament length, intensity ratio, hyphal diameter and distance between branching

We measured filament length of branch points by merging both Cdc11a and Cdc11b channels using ImageJ. Measurements of filament lengths were made manually. For ratiometric analysis, images were background subtracted (in both channels individually). Summed fluorescence intensity was exacted at each of these sites. The summed fluorescence intensity sum of the Cdc11a channel was then divided by the summed fluorescence intensity of the Cdc11b channel. Measurements from all structures was plotted using PlotsofData (Postma and Goedhart, 2019). Hyphal diameters and the distance between branches were measured from DIC images of various strains using ImageJ.

## Tables

### Strains used in this study

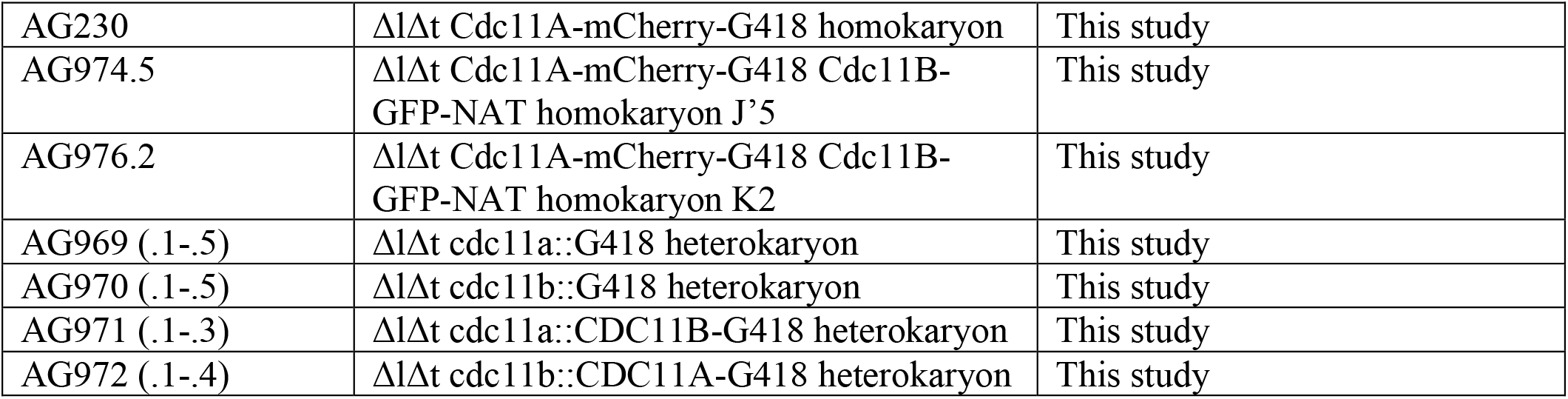

### Plasmids used in this study

|  |  |  |
| --- | --- | --- |
| AGB048 | pAGT211 pUC19 mCherry-NAT | Gift from Basel |
| AGB125 | pRS416 AgCDC11A | Basel |
| AGB141 | pRS416 AgCDC11A-mCherry-NAT | This study |
| AGB005 | pAGT141 pG3GFP | Basel |
| AGB124 | pRS416 AgCDC11B | Basel |
| AGB1320 | pRS416 AgCdc11b-GFP-G418 | This study |
| AGB1369 | pcdc11a::G418 | This study |
| AGB1370 | pcdc11b::G418 | This study |
| AGB1374 | pUC19 cdc11a::CDC11B-GFP-NEO | This study |
| AGB1352 | pUC19 cdc11b::CDC11A-GFP-NEO | This study |
| AGB 119 | pACYC AgCDC3/AgCDC11A | This study |
| AGB827 | pETDuet ScCDC12/AgCDC10 | This study |

### Oligonucleotides used in this study

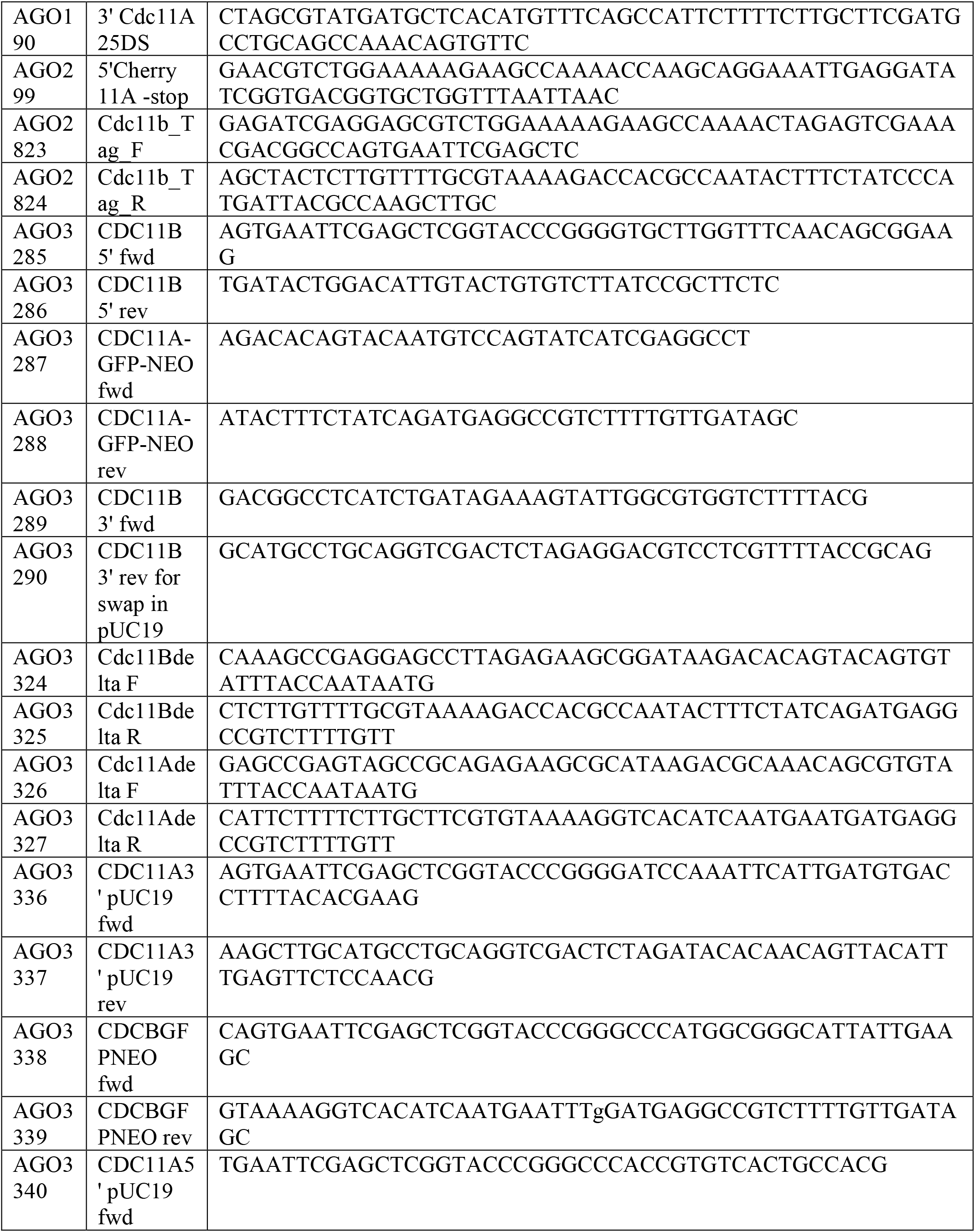
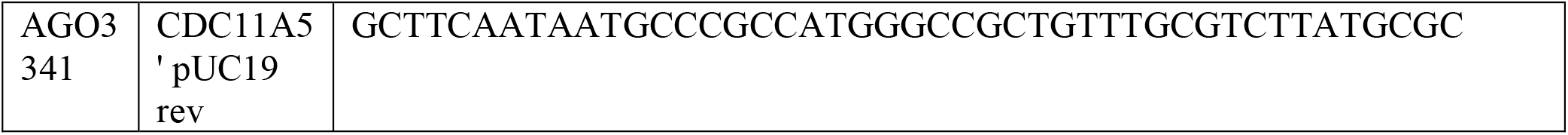

## Supplemental material

**Figure S1.**
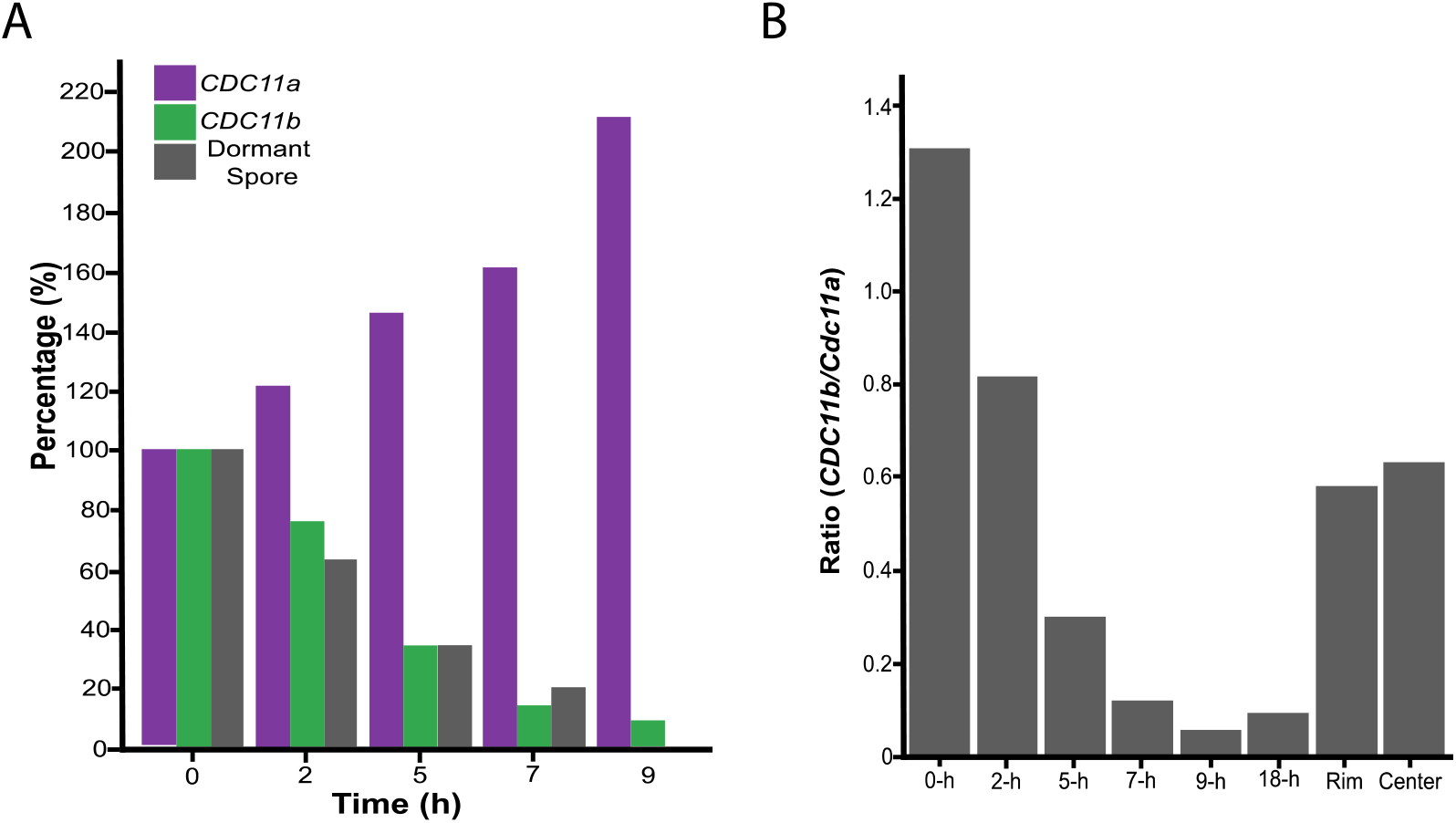
*CDC1b* transcription is regulated over time. (A) *CDC11b* transcription is repressed in early time points in liquid media. *CDC11b* transcript is relatively abundant in the spores (0-h), and repressed after germination. (B) Spores contain a higher ratio of *CDC11B/CDC11a* transcript. After germination, *CDC11a* transcripts are more abundant. In mature mycelia, the ratio of *CDC11a*:*CDC11b* is closed to 2:1

**Figure S2.**
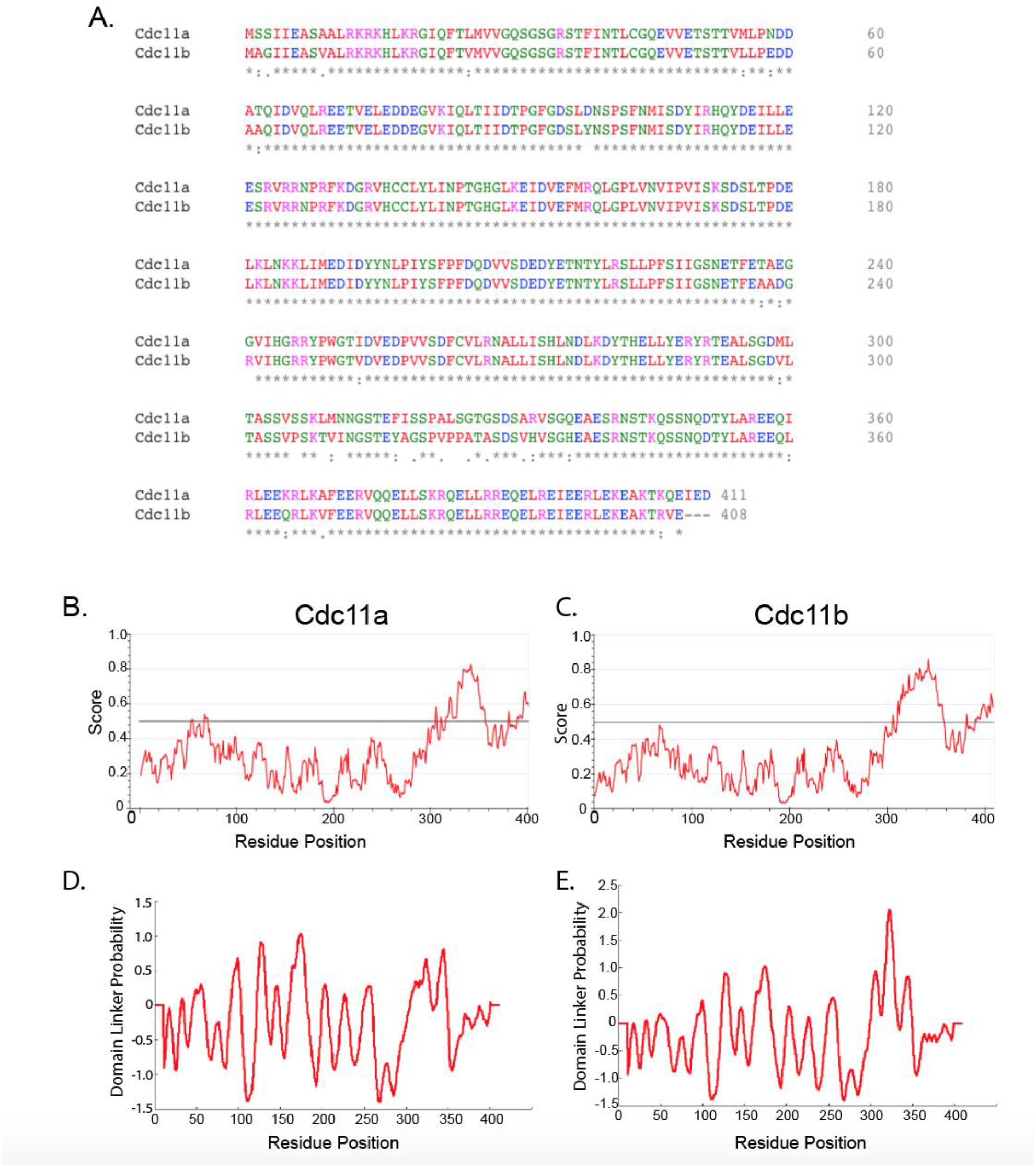
The highest degree of sequence variation between Cdc11a and Cdc11b is within the C-terminal extension. (A) Primary sequence alignment of Cdc11a and Cdc11b. (B,C) IUPRED output of predicted intrinsically disordered regions within Cdc11a (panel 1) and Cdc11b (panel 2) polypeptides. (D,E) Loop-length-dependent support vector machine prediction of protein domain linkers in Cdc11a and Cdc11b polypeptides, respectively.

## Acknowledgements

We would like to thank members of the Gladfelter lab for lively and stimulating discussions around septins. Kevin would also like to thank Joanne Ekena for having perseverance, patience, grace, and the heart of a champion to make strains. This work was supported by NSF MCB 1615138 and MCB-2016022, and an HHMI Faculty Scholars award to ASG, NIH. KSC was supported in part by a grant from the National Institute of General Medical Sciences under award T32 GM119999. JMV-M was supported by NIH Training Grant 5T32AI052080-14.

## References

Auxier, B. et al.. (2019) ‘Diversity of opisthokont septin proteins reveals structural constraints and conserved motifs 06 Biological Sciences 0604 Genetics’, BMC Evolutionary Biology. doi: 10.1186/s12862-018-1297-8.

Ayad-Durieux, Y. et al.. (2000) ‘A PAK-like protein kinase is required for maturation of young hyphae and septation in the filamentous ascomycete Ashbya gossypii’, Journal of Cell Science.

Bertin, A. et al.. (2008) ‘Saccharomyces cerevisiae septins: Supramolecular organization of heterooligomers and the mechanism of filament assembly’, Proceedings of the National Academy of Sciences. doi: 10.1073/pnas.0803330105.

Bertin, A. et al.. (2010) ‘Phosphatidylinositol-4,5-bisphosphate Promotes Budding Yeast Septin Filament Assembly and Organization’, Journal of Molecular Biology. doi: 10.1016/j.jmb.2010.10.002.

Bridges, A. A. et al.. (2014) ‘Septin assemblies form by diffusion-driven annealing on membranes’, Proceedings of the National Academy of Sciences. doi: 10.1073/pnas.1314138111.

Bridges, A. A. et al.. (2016) ‘Micron-scale plasma membrane curvature is recognized by the septin cytoskeleton’, Journal of Cell Biology. doi: 10.1083/jcb.201512029.

Bridges, A. A. and Gladfelter, A. S. (2016) ‘In vitro reconstitution of septin assemblies on supported lipid bilayers’, Methods in Cell Biology. doi: 10.1016/bs.mcb.2016.03.025.

Cannon, K. S. et al.. (2019) ‘An amphipathic helix enables septins to sense micrometer-scale membrane curvature’, Journal of Cell Biology. doi: 10.1083/jcb.201807211.

DeMarini, D. J. et al.. (1997) ‘A septin-based hierarchy of proteins required for localized deposition of chitin in the Saccharomyces cerevisiae cell wall’, Journal of Cell Biology. doi: 10.1083/jcb.139.1.75.

DeMay, B. S. et al.. (2009) ‘Regulation of distinct septin rings in a single cell by Elm1p and Gin4p kinases.’, Molecular biology of the cell. doi: 10.1091/mbc.E08-12-1169.

Field, C. M. et al.. (1996) ‘A purified Drosophila septin complex forms filaments and exhibits GTPase activity’, Journal of Cell Biology. doi: 10.1083/jcb.133.3.605.

Garcia, G. et al.. (2011) ‘Subunit-dependent modulation of septin assembly: Budding yeast septin Shs1 promotes ring and gauze formation’, Journal of Cell Biology. doi: 10.1083/jcb.201107123.

Gattiker, A. et al.. (2007) ‘Ashbya Genome Database 3.0: A cross-species genome and transcriptome browser for yeast biologists’, BMC Genomics. doi: 10.1186/1471-2164-8-9.

Gilden, J. and Krummel, M. F. (2010) ‘Control of cortical rigidity by the cytoskeleton: Emerging roles for septins’, Cytoskeleton. doi: 10.1002/cm.20461.

Gladfelter, A. S. et al.. (2005) ‘Interplay between septin organization, cell cycle and cell shape in yeast.’, Journal of cell science. doi: 10.1242/jcs.02286.

Graham, J. S. et al.. (2014) ‘Multi-platform compatible software for analysis of polymer bending mechanics’, PLoS ONE. doi: 10.1371/journal.pone.0094766.g001.

Hartwell, L. H. et al.. (1974) ‘Genetic control of the cell division cycle in yeast’, Science. doi: 10.1126/science.183.4120.46.

Helfer, H. and Gladfelter, A. S. (2006) ‘AgSwe1p regulates mitosis in response to morphogenesis and nutrients in multinucleated Ashbya gossypii cells’, Molecular Biology of the Cell. doi: 10.1091/mbc.E06-03-0215.

Hilary Russell, S. E. and Hall, P. A. (2011) ‘Septin genomics: A road less travelled’, in Biological Chemistry. doi: 10.1515/BC.2011.079.

Howell, A. S. et al.. (2009) ‘Singularity in Polarization: Rewiring Yeast Cells to Make Two Buds’, Cell. doi: 10.1016/j.cell.2009.10.024.

Kelley, J. B. et al.. (2015) ‘RGS proteins and septins cooperate to promote chemotropism by regulating polar cap mobility’, Current Biology. doi: 10.1016/j.cub.2014.11.047.

Kelley, L. A. et al.. (2015) ‘The Phyre2 web portal for protein modeling, prediction and analysis’, Nature Protocols. doi: 10.1038/nprot.2015.053.

Khan, A., Newby, J. and Gladfelter, A. S. (2018) ‘Control of septin filament flexibility and bundling by subunit composition and nucleotide interactions’, Molecular Biology of the Cell. doi: 10.1091/mbc.E17-10-0608.

Knechtle, P., Dietrich, F. and Philippsen, P. (2003) ‘Maximal polar growth potential depends on the polarisome component AgSpa2 in the filamentous fungus Ashbya gossypii’, Molecular Biology of the Cell. doi: 10.1091/mbc.E03-03-0167.

Köhli, M. et al.. (2008) ‘Growth-speed-correlated localization of exocyst and polarisome components in growth zones of Ashbya gossypii hyphal tips’, Journal of Cell Science. doi: 10.1242/jcs.033852.

Luedeke, C. et al.. (2005) ‘Septin-dependent compartmentalization of the endoplasmic reticulum during yeast polarized growth’, Journal of Cell Biology. doi: 10.1083/jcb.200412143.

Mela, A. and Momany, M. (2021) ‘Septins coordinate cell wall integrity and lipid metabolism in a sphingolipid-dependent process’, Journal of Cell Science. doi: 10.1242/jcs.258336.

Ong, K. et al.. (2014) ‘Architecture and dynamic remodelling of the septin cytoskeleton during the cell cycle’, Nature Communications. doi: 10.1038/ncomms6698.

Pan, F., Malmberg, R. L. and Momany, M. (2007) ‘Analysis of septins across kingdoms reveals orthology and new motifs’, BMC Evolutionary Biology. doi: 10.1186/1471-2148-7-103.

Postma, M. and Goedhart, J. (2019) ‘Plotsofdata—a web app for visualizing data together with their summaries’, PLoS Biology. doi: 10.1371/journal.pbio.3000202.

Romberg, L., Simon, M. and Erickson, H. p. (2001) ‘Polymerization of FtsZ, a bacterial homolog of tubulin. Is assembly cooperative?’, Journal of Biological Chemistry. doi: 10.1074/jbc.M009033200.

Schmitz, H. P. et al.. (2006) ‘From function to shape: A novel role of a formin in morphogenesis of the fungus Ashbya gossypii’, Molecular Biology of the Cell. doi: 10.1091/mbc.E05-06-0479.

Seiler, S. and Plamann, M. (2003) ‘The Genetic Basis of Cellular Morphogenesis in the Filamentous Fungus Neurospora crassa’, Molecular Biology of the Cell. doi: 10.1091/mbc.E02-07-0433.

Sellin, M. E., Stenmark, S. and Gullberg, M. (2014) ‘Cell type-specific expression of SEPT3- homology subgroup members controls the subunit number of heteromeric septin complexes’, Molecular Biology of the Cell. doi: 10.1091/mbc.E13-09-0553.

Shen, X. X. et al.. (2018) ‘Tempo and Mode of Genome Evolution in the Budding Yeast Subphylum’, Cell. doi: 10.1016/j.cell.2018.10.023.

Skillman, K. M. et al.. (2013) ‘The unusual dynamics of parasite actin result from isodesmic polymerization’, Nature Communications. doi: 10.1038/ncomms3285.

Steenwyk, J. L. and Rokas, A. (2019) ‘Treehouse: A user-friendly application to obtain subtrees from large phylogenies’, BMC Research Notes. doi: 10.1186/s13104-019-4577-5.

Steger, G. (1998) ‘An unbiased detector of curvilinear structures’, IEEE Transactions on Pattern Analysis and Machine Intelligence. doi: 10.1109/34.659930.

Woods, B. L. et al.. (2021) ‘Biophysical properties governing septin assembly’, bioRxiv.

Yamada, S. et al.. (2016) ‘Septin Interferes with the Temperature-Dependent Domain Formation and Disappearance of Lipid Bilayer Membranes’, Langmuir. doi: 10.1021/acs.langmuir.6b03452.

